# Rapid single-molecule characterisation of nucleic-acid enzymes

**DOI:** 10.1101/2022.03.03.482895

**Authors:** Stefan H. Mueller, Lucy J. Fitschen, Afnan Shirbini, Samir M. Hamdan, Lisanne M. Spenkelink, Antoine M. van Oijen

## Abstract

The activity of enzymes is traditionally characterised through bulk-phase biochemical methods that only report on population averages. Single-molecule methods are advantageous in elucidating kinetic and population heterogeneity but are often complicated, time consuming, and lacking statistical power. We present a highly generalisable and high-throughput single-molecule assay to rapidly characterise proteins involved in DNA metabolism. The assay exclusively relies on changes in total fluorescence intensity of surface-immobilised DNA templates as a result of DNA synthesis, unwinding or digestion. Combined with an automated data-analysis pipeline, our method provides enzymatic activity data of thousands of molecules in less than an hour. We demonstrate our method by characterising three fundamentally different nucleic-acid enzyme activities: digestion by the phage *λ* exonuclease, synthesis by the phage *Phi29* polymerase, and unwinding by the *E. coli* UvrD helicase. We observe a previously unknown activity of the UvrD helicase to remove proteins tightly bound to the ends of DNA.

## MAIN

Maintenance of DNA, involving replication, repair, and recombination, requires many different enzymes with a range of different activities. Development of information-rich biochemical assays that report on these activities is an important step towards our understanding of their molecular mechanisms in disease pathways such as anti-microbial resistance [1] and cancer [2]. Additionally, characterisation of nucleic-acid enzymes plays an important role in the development of methods such as gene amplification and DNA sequencing, widely used not only in molecular biochemistry, but also forensics, diagnostics [3] and palaeontology [4]. Traditionally, the activity of DNA-modifying enzymes is characterised through ensemble-averaging biochemical methods, such as gel electrophoresis and fluorimetry. These methods have the drawback of averaging over large ensembles of molecules and, therefore, provide no access to information on subpopulations, dynamic molecular mechanisms and intermediate states. However, knowledge of these properties is often crucial to a full understanding of the molecular processes underlying DNA metabolism and the enzymes involved.

To describe such properties, researchers have developed techniques to observe single molecules in real time. These methods often rely on imaging fluorescent tags or manipulating molecules using optical tweezers [5]. In recent years these techniques have revealed unexpected dynamics [6]-[9] and quantitatively characterised interactions on the molecular scale [10], [11]. While these techniques have yielded new insights into molecular properties of enzymes and protein dynamics, a major disadvantage of single-molecule approaches is the time-consuming and complex nature of the experiments and data analysis needed to acquire statistically significant data. Because of these challenges, single-molecule studies are difficult to reproduce by other researchers and the statistical power of many studies is comparatively small. Here, we describe a single-molecule assay that can be used to characterise any enzyme that catalyses the conversion between double-stranded DNA (dsDNA) and single-stranded DNA (ssDNA). The assay provides kinetic information on large numbers of molecules in one experiment and is simple to implement relative to existing single-molecule experiments. By using fluorescent probes that selectively stain ssDNA or dsDNA and by monitoring fluorescence intensity changes of surface-immobilised, randomly-coiled DNA templates, we can visualise the conversion of dsDNA to ssDNA in real time for hundreds of molecules simultaneously.

As a proof of principle, we characterise three enzymes with different functions. We visualise exonucleolytic degradation of the DNA template catalysed by phage λ exonuclease (λ exo) (Fig. 1a), strand-displacement synthesis by the phage Phi29 DNA polymerase (Phi29 DNAp) (Fig. 1b), and unwinding of DNA by the *E. Coli* UvrD helicase (Fig. 1c). We report rate constants and distributions, determined by characterising thousands of single-molecule reactions for each of the three enzymes. The statistical power of our study greatly exceeds that of previous single-molecule studies of these enzymes ([12]-[15], yet our assay is comparatively easy to implement and due to a highly automated data-analysis pipeline, less time-consuming than previously described methods. Using this assay we observed the removal of protein roadblocks from DNA ends by the UvrD helicase — an activity that was previously unknown.

**Fig. 1:**
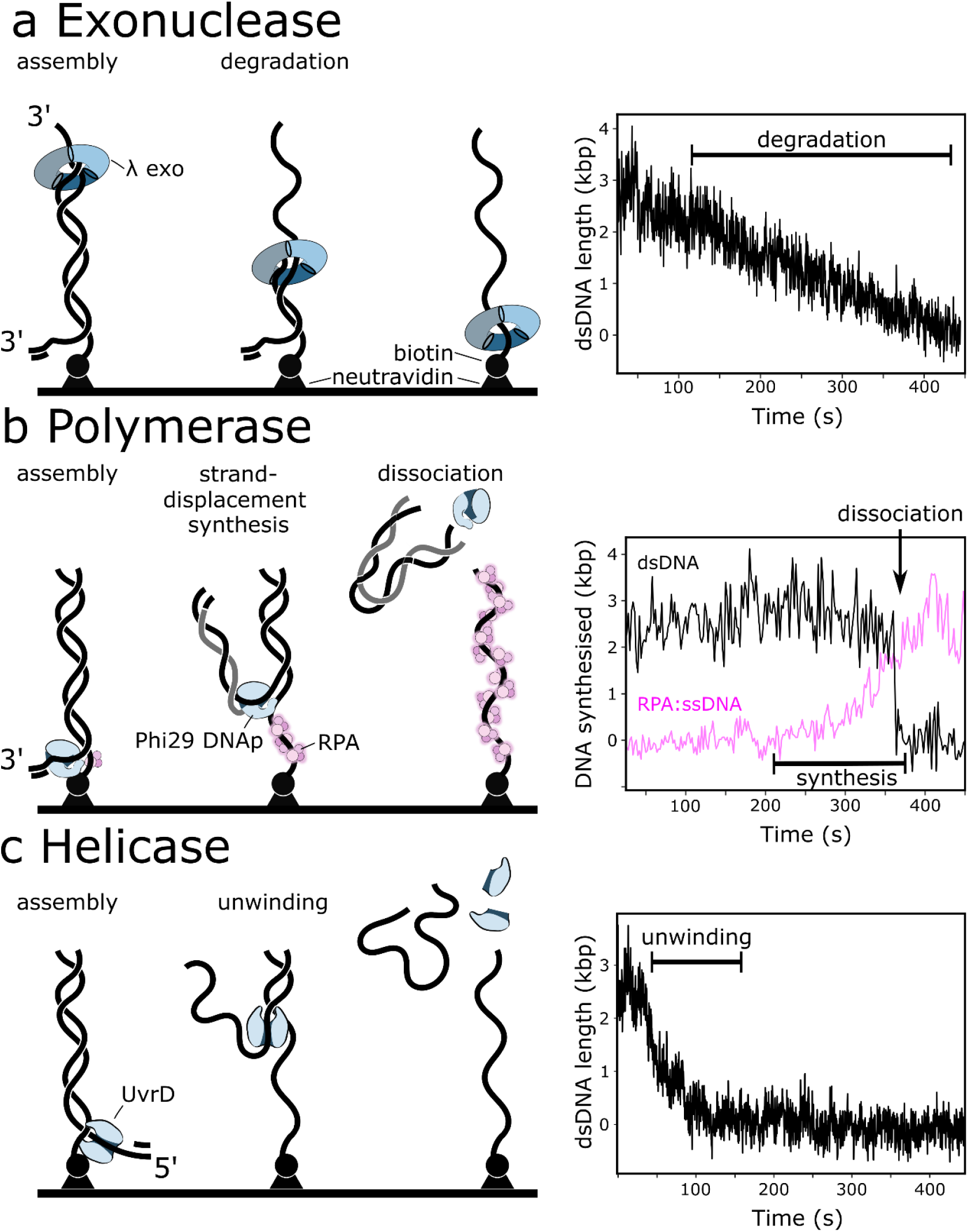
Schematic representation of the assay and example data. **(a)** Trimeric exonuclease lambda loads on free 3’ ends and converts dsDNA to ssDNA (left). This digestion leads to a decrease in measured DNA stain intensity (right). **(b)** Phi29 DNAp mediated strand-displacement synthesis (left). Phi29 DNAp binds the primed 3’ end of the template. In the presence of dNTPs the template strand becomes single stranded, as the newly-synthesised daughter strand is displaced and eventually dissociates from the surface and therefore becomes invisible in TIRF-microscopy. This displacement leads to the instantaneous drop in DNA stain intensity (right; black line). In parallel, we use fluorescently-labeled RPA to visualise the increasing amount of exposed ssDNA (magenta line). **(c)** UvrD helicase assembles at the available 5’ end (left). In the presence of ATP dsDNA is converted to ssDNA, leading to a decrease in DNA stain intensity (right).

## RESULTS

### Single-molecule characterisation of λ exonuclease activity

Our assay uses easily-constructed 2.6-kb linear dsDNA templates. These templates have a biotin on one end to allow for attachment to the surface of a microfluidic flow chamber through biotin-neutravidin binding. After surface immobolisation of the templates, we stain the dsDNA by introducing the DNA-intercalating dye SYTOX Orange into the flow cell. Finally, we initiate the enzymatic reaction by adding the enzyme and required cofactors. The activity of any enzyme that alters the amount of dsDNA can be monitored by measuring SYTOX Orange fluorescence intensity.

The trimeric λ exo catalyses the removal of nucleotides from linear or nicked dsDNA in the 5’ to 3’ direction. During degradation of DNA, the enzyme encircles both strands [16], [17]. On our template, λ exo loads at the free, non-tethered end and subsequently converts the dsDNA to ssDNA by digesting nucleotides from the 5’ end (Fig. 1a). As dsDNA is converted to ssDNA staining by SYTOX Orange becomes much weaker. We can therefore monitor the digestion of dsDNA in real-time by integrating the DNA-stain intensity for each individual molecule over time (see Fig. 2 a,b). Over a total of six experiments we record trajectories of over 2500 individual molecules. Our method is sensitive to complex stochastic kinetics of individual molecules, such as pausing (see Fig. 2a, middle trajectory). However, the vast majority of trajectories shows very uniform and linear behaviour. We use piecewise linear fits to determine the digestion rate for individual molecules. We find rate distributions with means of 10.9±7.1 and 21.4±6.2 (mean ± standard deviation (STD)) for reactions at 25°C or 35°C respectively (see Fig. 2c), consistent with previously measured values [18] [12], [19]. The large standard deviation of the observed distributions highlights the presence of intermolecular disorder. Such effects have previously been studied by single-molecule techniques, but typically with much smaller sample sizes [12], [20], [21].

**Fig. 2.**
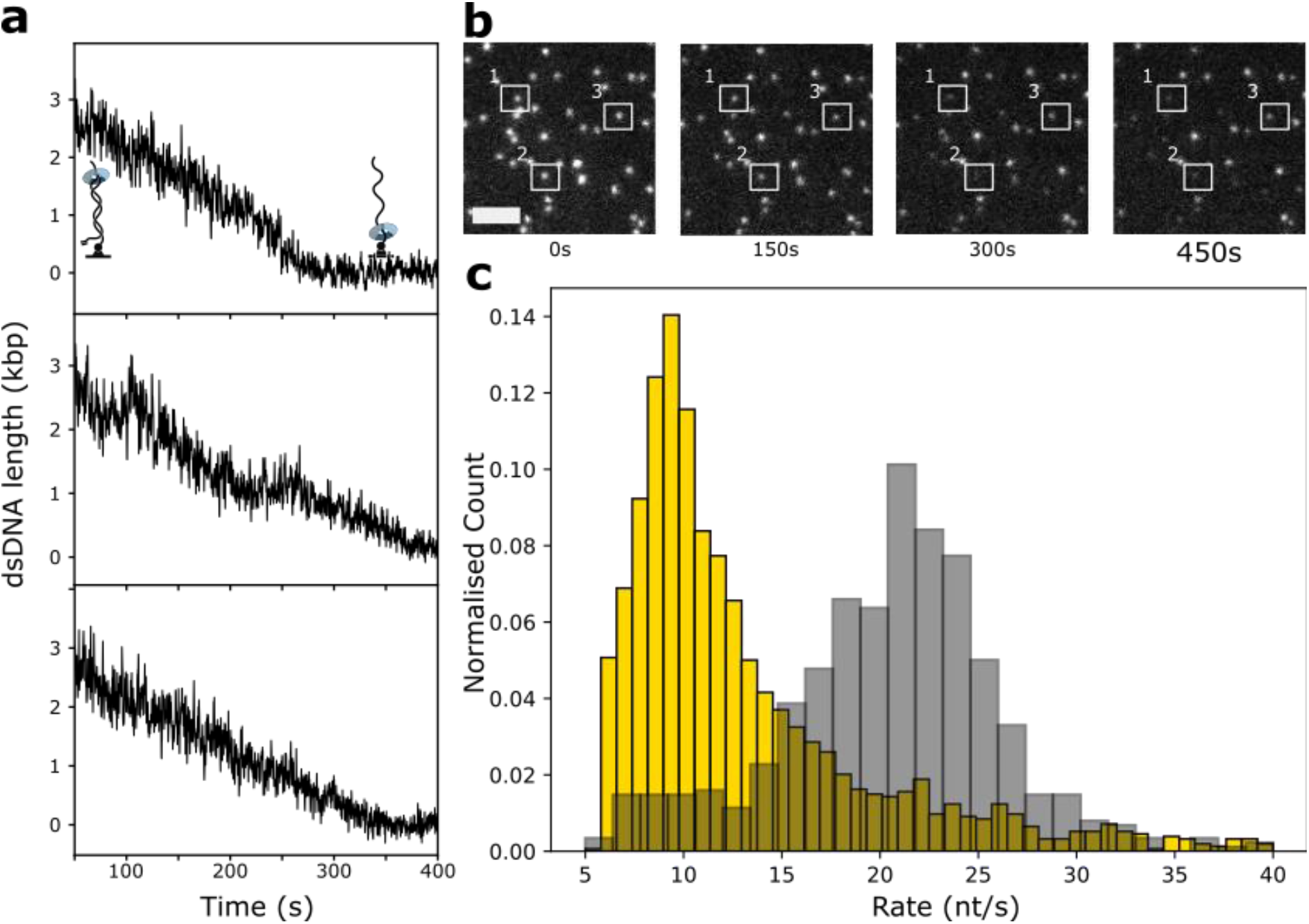
λ exonuclease. **(a)** Single-molecule trajectories of DNA degradation by λ exonuclease. **(b)** Still frames from a recorded movie with the white rectangles numbered 1-3 as a visual guide and marking identical molecules over time, showing their decrease in DNA-stain intensity. Scale bar: 5 μm. **(c)** Rate distributions of DNA degradation by λ exonuclease at 25°C(yellow, n = 1957 molecules) and 35 °C (grey, n = 648 molecules).

### Single-molecule characterisation of strand-displacement synthesis by the Phi29 DNA polymerase and ssDNA-binding properties of RPA

During strand-displacement synthesis by Phi29 DNAp, the net amount of dsDNA stays constant until the template strand is fully replicated and the daughter strand dissociates from the template (see Fig. 2a). This dissociation is visible as a sudden drop in DNA-stain intensity. To visualise the kinetics of DNA synthesis, we additionally introduce fluorescently-labelled *S. cerevisiae* replication protein A (RPA), a single-stranded DNA-binding protein with very high affinity for ssDNA. Furthermore, while free RPA is present in solution, bound RPA exchanges rapidly [22] [23]. This effect mitigates photobleaching and makes RPA a good marker for ssDNA (see supplementary Fig 1). For every synthesised nucleotide on the daughter strand, one nucleotide of ssDNA is left behind on the surface-tethered strand. Therefore, the change in RPA signal over time corresponds to the replication rate by Phi29 DNAp, knowing that RPA binds ssDNA faster than new dsDNA is synthesised [24]. Fig. 3c shows data from our experiments. Surprisingly, most single-molecule trajectories seemed to exhibit a non-linear dependence between RPA signal and time. However, for individual traces this behaviour is difficult to distinguish from statistical noise and pausing kinetics. To increase our signal-to-noise ratio we synchronised all trajectories to the time of dissociation of the daughter strand, which is an event that is easy to identify. Subsequent averaging across many single-molecule traces yields a synchronised average trajectory that does not suffer from the same caveats as ensemble-averaging methods and contains information on the underlying kinetics [25], [26]. Indeed, the post-synchronised average trajectory clearly exhibits non-linear behaviour (see Fig. 3d). As a control, to prove that this is not an artefact of our analysis, we synchronised trajectories from the previous experiments on λ exonuclease (see supplementary Fig 2). Unlike for Phi29 DNAp, the synchronised trajectory of λ exonuclease is linear. The observed non-linear Phi29 DNAp dynamics become more obvious for higher dNTP concentrations, i.e. at higher replication rates (see Fig. 3d). This observation suggests that our initial assumption, that RPA binding kinetics are faster than DNA synthesis, is not generally true. Our data is well described by single-exponential functions, with a *K_M_* of Phi29 DNAp for dNTPs of (8±3) μM (mean±SEM) and a maximum synthesis rate of 160±25 bp·s^-1^ (mean±SEM, see Fig. 3d). Our *K_M_* value is about four times lower than previously reported values and our *v_max_* value is consistent with previous measurements [27]. The fact that we find lower *K_M_* values than expected is in agreement with the hypothesis that for high dNTP concentrations, the *observed* kinetics are limited by RPA binding. This is also confirmed by the fact that our estimated *v_max_* is in good agreement with the literature, since the maximum increase in RPA signal is still limited by strand-displacement synthesis by Phi29 DNAp.

**Fig. 3.**
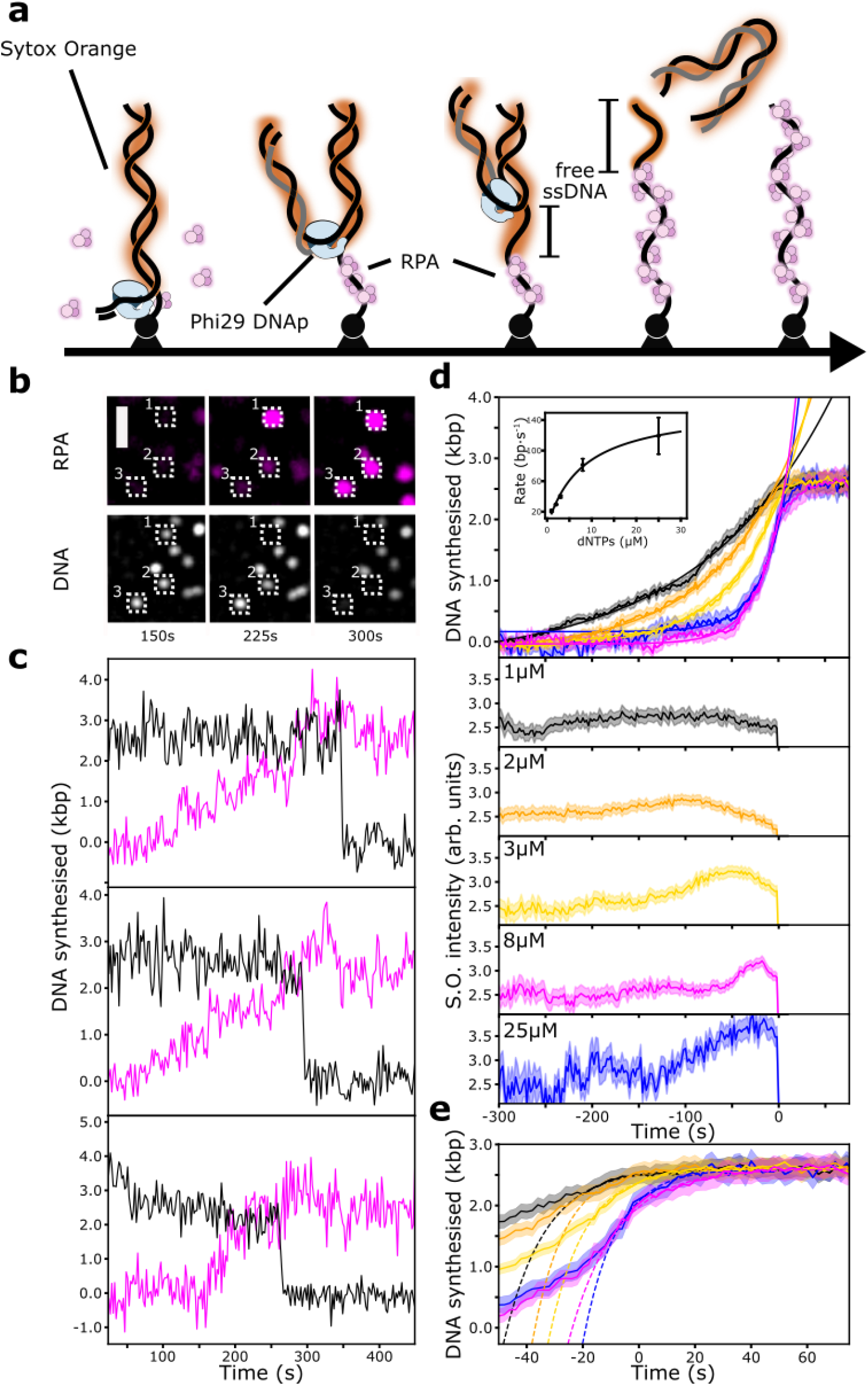
Single-molecule characterisation of Phi29 DNAp. **(a)** Schematic representation of the assay. First, Phi29 DNAp assembles on the primed DNA end. As ssDNA is created, free ssDNA accumulates before it is bound by RPA. Eventually the daughter strand dissociates, leaving behind ssDNA bound by RPA. **(b)** Montage of a TIRFM movie of strand-displacement synthesis over 300 s. The RPA signal (magenta, top) increases in intensity, while the dsDNA signal (white, bottom) stays approximately constant until it finally disappears. Scale bar: 2 μm. **(c)** Three individual single-molecule trajectories of strand-displacement synthesis. (dsDNA signal: black, ssDNA/RPA signal: magenta). **(d)** Post-synchronised average trajectories at dNTP concentrations (each) of 1 μM (black, n=101 molecules), 2 μM (orange, n=153 molecules), 3 μM (yellow, n=134 molecules), 8 μM (magenta, n=127 molecules) and 25 μM (blue, n=58 molecules). The shaded area around the lines depicts the standard error of the mean (SEM). The top graph shows the RPA signal, normalised to the length of the template DNA (2620 bp). Solid smooth lines are exponential fits (see methods) of the data, restricted to time values smaller than −10 seconds. Rate constants were obtained by multiplying with the length of the template DNA. The average rate in nt·s^-1^ was determined from three independent experiments and is shown in the inset. The error bars show the standard error of the mean. The solid line in the inset is a fit to the Michaelis-Menten equation (see methods) yielding: K_M_ = 8 ± 3 μM, v_max_ = 158 ± 25 bp·s^-1^. The bottom five graphs show the synchronised DNA-stain signals over time. **(e)** The increase in RPA signal prior to dissociation of the daughter strand. The dashed lines are fits to first-order rate equations (see methods). The extracted association constant from a total of 15 datasets is 11.1 ± 0.9 nt·nM^-1^·s^-1^(mean ± SEM).

To gain information on RPA binding kinetics within our system we examine the increase in RPA fluorescence signal immediately before the dissociation of the daughter strand (see Fig. 4c). In this regime RPA binding is no longer limited by Phi29 DNAp activity. The observed data are well described by first-order binding kinetics and yield a bimolecular association rate constant *k_on_* of 11.1±0.9 nt·nM^-1^·s^-1^ (mean±SEM), consistent with previously reported values [24], [28]. A first-order kinetic model for RPA binding implies that the speed of binding at any given time depends on the number of binding partners available (see methods). We therefore hypothesise that in the very beginning of the reaction almost no free ssDNA is present, and RPA binding is therefore slow. As replication proceeds ssDNA is generated. As more binding sites become available, RPA binding becomes faster, the amount of ssDNA decreases again, and binding slowly converges to saturation as the daughter strand dissociates (see Fig. 4a). This picture implies a fluctuation of the amount of free ssDNA, not bound by RPA but very weakly stained by Sytox Orange. Since ssDNA staining is much less efficient than dsDNA, such a minor increase is not visible in the individual single-molecule trajectories. However, the post-synchronised trajectories of the S.O. signal (see Fig. 4d) indeed show a clear fluctuation in intensity. Our data indicates that RPA binding is stimulated by the amount of free ssDNA, and that RPA displaces Sytox Orange from ssDNA.

**Fig. 4.**
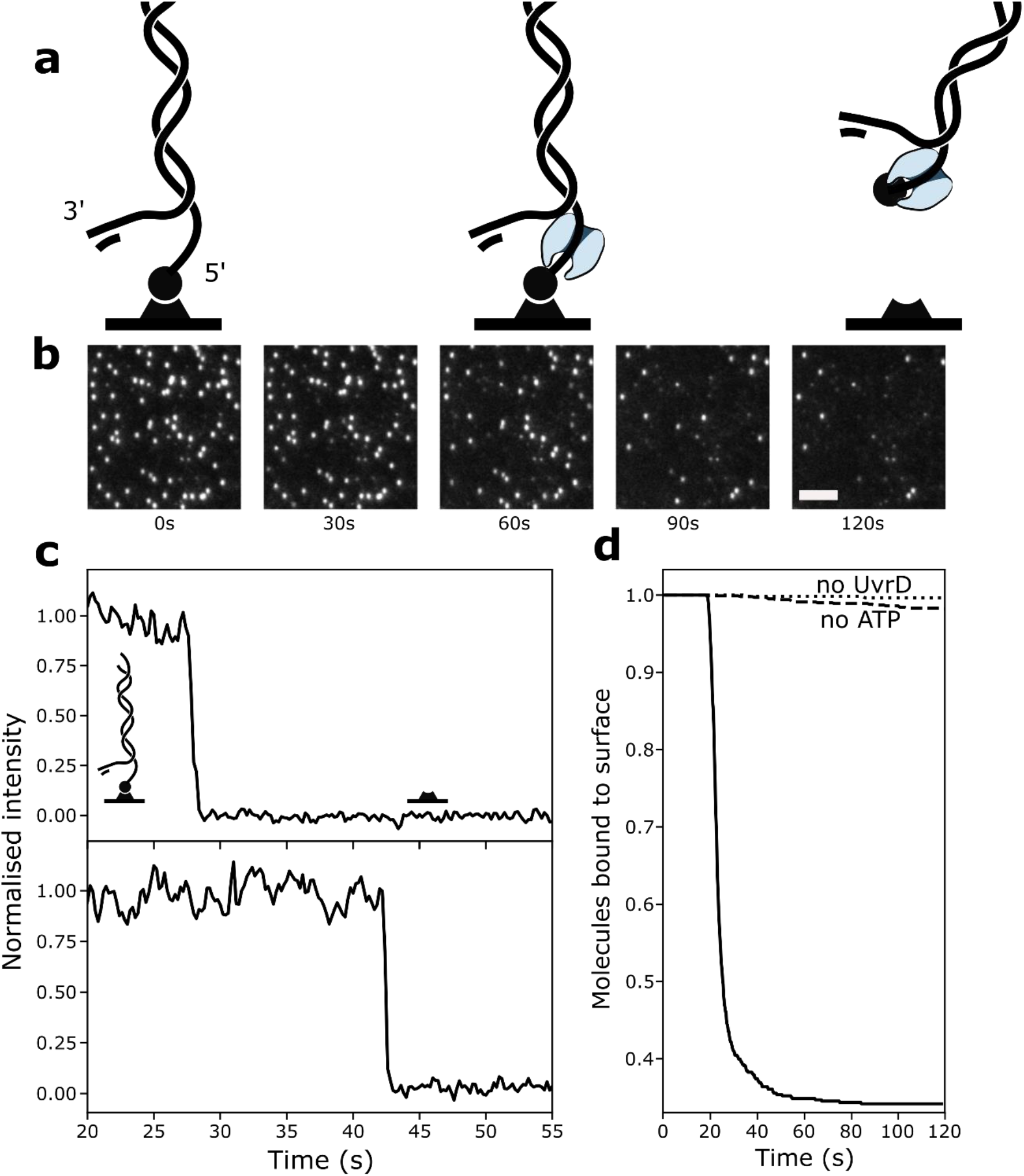
UvrD disrupts biotin-neutravidin interaction. **(a)** Schematic representation of UvrD assembly next to the tethered 5’ end and subsequent dissociation of the DNA substrate. **(b)** Montage of a recorded movie. The bright spots correspond to individual DNA molecules disappearing over time. **(b)** Integrated fluorescence intensities over time from the movie shown in (a). **(c)** The drop in intensity corresponds to the removal of the neutravidin from the biotinylated DNA end. **(d)** Fraction of DNA substrates still bound to the surface over time, in presence of UvrD and ATP (solid line, n=588 molecules), absence of UvrD or ATP respectively (dotted line, n=1004, dashed line n = 583 molecules).

### Characterisation of the *E. coli* UvrD helicase

Next, we sought to test if our assay is suitable for the study of helicases. Helicases are one of the biggest families of proteins, present in all domains of life. As an example we characterise the *E. coli* UvrD helicase, a member of the SF1 family of helicases. Apart from unwinding DNA in 3’ to 5’ direction in its dimeric form, it is also involved in methyl-directed mismatch repair and acts as an antirecombinase by removing recA filaments from ssDNA [29]-[31]. The monomeric form of UvrD processively translocates on ssDNA [15].

At first, we wanted to study ATP-dependent unwinding by the UvrD helicase on the previously used 2.6-kb forked template. We expected unwinding of DNA, and therefore a continuous decrease in the fluorescence intensity of S.O. stained template DNA, as UvrD unwinds the substrate, potentially from both ends. Surprisingly, instead of the expected continuous decrease in intensity, we observe a discrete drop in fluorescence intensity, i.e., diffraction limited spots simply disappear (see Fig. 5a). This observation indicates dissociation of the full template from the cover slide surface rather than DNA unwinding. Together with control reactions lacking either ATP or using inactivated UvrD, this shows that the UvrD helicase can actively remove the neutravidin bound to the 5’-DNA end. We The trajectories still show discrete fluorescence drops within one frame, suggesting a dissociative process that is completed within 30 ms. UvrD loading on the free 3’-DNA end and unwinding DNA towards the surface within 30 ms would correspond to a rate of 80,000 nt·s^-1^. Since this high rate would be in stark contradiction to the literature [14], [30], [32], [33], we conclude that UvrD is loading in close proximity to the surface and exhibits an enzymatic activity different to DNA unwinding.

**Fig. 5.**
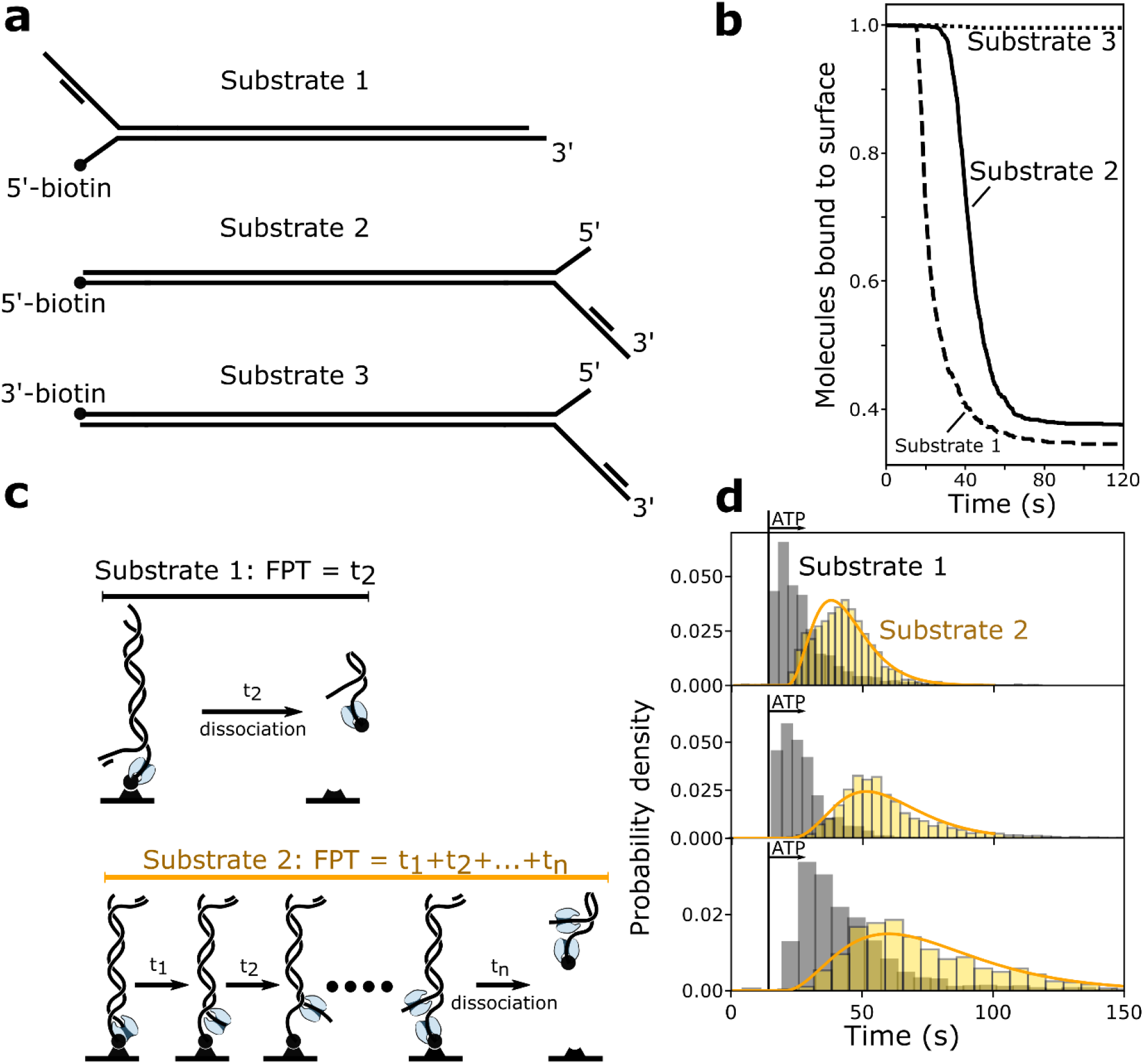
Substrate and ATP dependence of UvrD-mediated removal of neutravidin. **(a)** Schematic overview of DNA substrates. Substrate 1: 5’ biotin with ssDNA overhang, Substrate 2:5’ biotin on blunt end, Substrate 3: 3’ biotin on blunt end. **(b)** Fraction of molecules bound to the coverslide surface over time for substrates 1-3 in presence of 50 μM ATP and 100 nM UvrD. **(c)** Principle of the First Passage Time (FPT) for a reaction without intermediate steps (top), resulting in exponentially distributed FPT distributions. For a reaction with n intermediate steps (bottom), the measured FPT equals the sum of intermediate steps. If all intermediate steps follow identical kinetics, the FPT distribution follows a gamma distribution. **(d)** FPT distributions for Substrates 1 (grey) and 2 (yellow) in presence of 100 nM UvrD and 50 μM ATP (top, Substrate 1: n=1171 molecules, Substrate2: n=2534 molecules), 25 μM ATP (middle, Substrate 1: n=817 molecules, Substrate2: n=2372 molecules) and 12.5 μM ATP (bottom, Substrate 1: n=973 molecules, Substrate2: n=1440 molecules). The vertical bar represents the time when ATP is first present in the flowcell. Note that the first bins of the histograms are likely underrepresentative of the distributions for two reasons: First, ATP concentration within the flowcell gradually increases to the given value and second, flow is operated manually which leads to an uncertainty of the time of ATP arrival in the order of seconds. The histograms contain data from three experiments each, with slightly different ATP arrival times. Orange lines are maximum-likelyhood estimations to gamma distributions, with rate parameter k and shape parameter n (50 μM: k=0.18s-1, n=4.2; 25 μM: k=0.12s-1, n=4.8; 12.5 μM: k=0.06s-1, n=3.4;).

Next, we wanted to understand if loading on ssDNA is required for the removal of neutravidin. To do so, we made two different versions of our previous DNA substrate (henceforth referred to as Substrate 1). First, we removed the ssDNA region adjacent to the tethered 5’-end (Substrate 2) to examine if displacement of protein blocks required UvrD assembly on ssDNA. Second, we placed the biotin on the 3’-DNA end (Substrate 3), to see if this activity has the same 3’-5’ directionality as unwinding and translocation on ssDNA (see Fig. 5a). DNA unwinding by UvrD was previously reported to be inefficient, if initiated from short 3’ overhangs or even blunt ends. Surprisingly, the removal of a neutravidin block is efficient, even from blunt ends (Fig. 5b, solid line). However, neutravidin bound to 3’-biotinilated DNA cannot be displaced by UvrD at all (see Fig. 5b, dotted line).

To gain more insight in the mechanism involved, we calculated first-passage time (FPT) distributions. FPT distributions are a powerful analysis tool, widely used to analyse and model stochastic processes, such as animal migration, the spread of COVID-19 virus particles and also helicase dynamics [34]-[36]. The FPT *t_n_* is the time from the start (addition of ATP) to the end of a reaction (dissociation of the DNA template) for an individual molecule (see Fig 5c). The distribution of FPTs conveys information on the number and rate constants of all rate-limiting steps during the reaction [37]. We preincubate Substrate 1 with UvrD and subsequently initiate the reaction by adding ATP. For Substrate 2, we observe a single-exponential FPT-distribution, a hallmark of the absence of intermediate steps (see Fig. 5c grey histograms). For Substrate 2, which lacks available ssDNA for UvrD to assemble close to the 5’ end, the FPTs are well described by a gamma distribution (see Fig 5b and c, yellow histrograms). This observation indicates the presence of multiple slow reaction intermediates required to remove the neutravidin [37]. Since the mean of the measured FPT distributions is much longer for Substrate 2 than for Substrate 1, we conclude that the rate-limiting steps in this case correspond to unwinding of the template from the blunt end (away from the surface), to subsequently allow for UvrD binding on the 5’ end next to the biotin. To obtain the number of reaction intermediates and corresponding rate-constants, we fit the data with gamma-distributions, as previously described [37] (see Fig. 5d and Methods). Our data suggest four intermediate steps, a number that does not vary with ATP concentration. Taken together with reported step sizes of unwinding by UvrD of 3–6 nucleotides [27], [38] our data indicates that unwinding of 12-24 nt is required for subsequent displacement of neutravidin.

Finally, we set out to observe DNA unwinding by UvrD. To do so, we utilise Substrate 3 (see Fig. 5a). The 3’-biotin prevents disruption of the biotin-neutravidin bond, while UvrD can load on the opposite end. The 60-nt 3’-dT tail provides a substrate for UvrD dimer assembly and initiation of DNA unwinding in presence of ATP. Unwinding by UvrD results in a gradual reduction of the DNA-stain intensity as dsDNA is converted to ssDNA. We find that UvrD is capable of unwinding the 2.6-kb template (Fig. 6 a,b). As before we use linear fits to determine a rate for each trajectory (see methods). We find a broad distribution, with a median of 29.5±28.3 bp·s^-1^ (median±STD, see Fig. 6c), consistent with values measured before in bulk and single-molecule studies [30], [38], [39]. However, to our knowledge, this is the first study reporting unwinding of long (>100 bp) DNA substrates, despite its potential importance during biological processes, such as methyl-directed mismatch repair, which can require more than 1000 bp to be unwound [40].

**Fig. 6.**
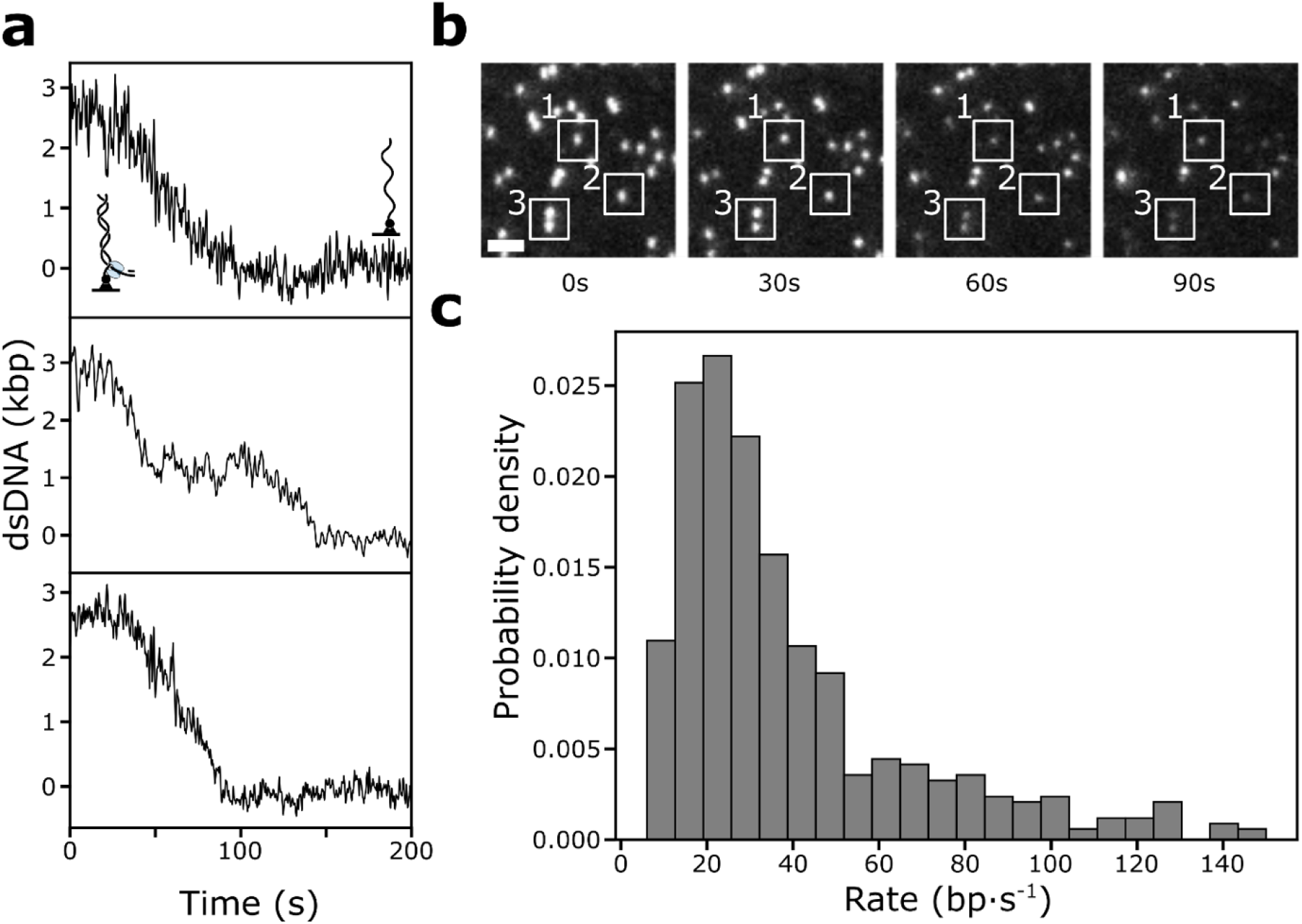
DNA unwinding by the UvrD helicase. **(a)** Single-molecule trajectories of DNA unwinding by UvrD. **(b)** Montage of TIRFM movie over 90s. Scale bar: 3 μm. The white rectangles serve as visual guide and mark individual molecules over time. **(c)** Rate distribution of DNA unwinding in the presence of 0.5 mM ATP at 25°C. n=452 molecules. The solid black line is a fit of the data to a bimodal distribution, the dashed lines represent the individual Gaussian components. Slow population: 13.2±2.5 bps^-1^ (mean±STD); fast population: 37.7±10.4 bp·s^-1^(mean±STD).

## Conclusions

We report a highly generalisable and high-throughput single-molecule assay with fully automated data analysis to study DNA-based enzymatic processes. This assay allows the extraction of features and kinetics otherwise hidden in the noise of single-molecule measurements. To demonstrate the strengths of this assay, we characterised DNA degradation, synthesis, and unwinding. Furthermore, we observe removal of a DNA-bound neutravidin by the UvrD helicase. Since proteins bound to DNA can present roadblocks to DNA replication in vivo [9], this new activity might have physiological relevance.

Reproducibility of fluorescence microscopy methods was previously identified as a major issue [41]. Quantitative fluorescence microscopy is inherently difficult to reproduce, due to the large number of factors involved. Fluorescence intensity varies dependent on the specific imaging apparatus, including the used light sources, as well as lenses and objectives and precise alignment thereof. Our assay produces data, in which mechanistic features are directly visible. This aspect allows for internal normalisation of fluorescence intensity and therefore circumvents this problem. Another factor of uncertainty in microscopy data is human bias during image analysis. We developed highly automated image analysis software for our assay, to minimise this problem.

Our method and analysis pipeline should be broadly applicable to measure the activity of any enzyme that converts dsDNA to ssDNA or *vice versa.* Furthermore, due to its high-throughput nature, the method has potential to be implemented in evolution or drug-screening studies.

## METHODS

### DNA template construction

As a starting material we used the 4kbf plasmid, a plasmid 4 kb in length and derived from pUC19, previously developed by Dr. Jacob Lewis. The plasmid was simultaneously digested with restriction endonucleases BsaI and BstXI (NEB). The resulting 2.6-kb fragment was separated from the 1.4-kb fragment and uncut plasmid by agarose gel purification (Promega Gel Wizzard Kit). A set of oligonucleotides that form a biotinylated and primed fork was ligated to one end of the fragment and the final product purified on a Sepharose-4B column as previously described [42]. The final product was stored at 4 °C. Full plasmid map and oligonucleotide sequences are described in the Supplementary Data section.

### Preparation of microfluidic flow cells

Flow chambers for microscopy were prepared as described before [7], [21], [43]. Briefly, cover slips (24 × 24 mm, Marienfeld) were functionalised with biotin-PEG (Laysan Bio). A polydimethylsiloxane (PDMS) block was made using soft-lithography methods and placed on top of the cover slips, creating a 1-mm wide and 0.1-mm high channel with a volume of 1 μL. Two stretches of polyethylene tubing (PE-60: 0.76-mm inlet diameter and 1.22-mm outer diameter, Walker Scientific) were inserted into the PDMS block at the entrance and exit of the channel to allow for buffer flow. Before the start of experiments, the flow channel was incubated with blocking buffer (50 mM Tris-HCl pH 7.6, 50 mM Potassium Chloride, 2% (v/v) Tween-20) to minimise nonspecific binding of DNA or proteins to the cover-slip surface. A syringe pump (Adelab Scientific) was used to introduce solutions to the flow cell.

### TIRF microscopy

The flow-cell device was mounted on an inverted total-internal reflection fluorescence (TIRF) microscope (Nikon Eclipse Ti-E), with an electrically heated stage (31°C; Okolab) and a 100x TIRF objective (NA = 1.49, oil, Nikon). Samples were illuminated using a 514-nm laser (Coherent, Sapphire 514-150 CW) at 1.6 mW cm^-2^ and a 647-nm laser (Coherent, Obis 647-100 CW) at 5.2 mW cm^-2^. The fluorescence signals were captured with an EMCCD camera (Hamamatsu C9100-13) through a dualband emission filter (TRF59907-EM, Chroma). For all measurements involving labelled RPA, samples were visualised at a frame rate of 0.5 frames per second with an exposure time of 400ms. For λ exo and UvrD reactions samples were visualised at a frame rate of 5 frames per second with an exposure time of 200ms, unless otherwise specified.

### λ exonuclease reactions

Firstly, 140 μL of 20 pM forked DNA template (Substrate 1) in Replication Buffer (25 mM Tris-HCl, pH 7.6, 10 mM magnesium acetate, 50 mM potassium glutamate, 40 μg/mL BSA, 0.1 mM EDTA, 5 mM dithiothreitol, and 0.0025% (v/v) Tween-20) in the presence of 150 nM Sytox Orange (Life Technologies) was loaded into the flowcell at a rate of 70 μL/min. After 1 minute or after a density of 0.3–0.7 molecules per μm^2^ on the surface was reached, 140 μL of Replication Buffer with 150 nM Sytox Orange and 20 nM RPA was loaded at a flow rate of 70 μL/min. Finally, 10 units of λ exonuclease (NEB) diluted in 80 μL of Replication Buffer supplemented with 150 nM Sytox Orange and 20 nM RPA were loaded into the flowcell at 70 μL/min.

### Strand-displacement reactions

First, the forked DNA template (20 pM in Replication Buffer) was loaded into the channel at a rate of 70 μL/min in the presence of 150 nM Sytox Orange (Life Technologies), allowing for direct visualisation. After 1 minute or after a density of 0.3–0.7 molecule per μm^2^ on the surface was reached, 80 μL of fluorescently labelled RPA (AF647-RPA, 20 nM in Replication Buffer supplemented with 150 nM Sytox Orange) was loaded at a rate of 70 μL/min. Purified RPA was a generous gift from Dr. Michael O’Donnell, fluorescent labelling of RPA was performed as previously described [7], [44]. Before the reaction was initiated, initial fluorescence intensities were recorded to determine the base line of RPA intensity at the fork. Next, 5 units of Phi29 DNAp (NEB) was loaded in the presence of 20 nM RPA and the specified concentration of dNTPs.

### UvrD cloning, expression and purification

The sequence-optimised UvrD gene tagged with 8xHis-tag at the N-terminus was cloned into pE-SUMO expression vector (Lifesensors Inc.) using Gibson reaction. The plasmid was then transformed by heat shock into BL21(DE3)pLysS *E. coli* competent cells. Four liters of 2YT media with a final concentration of 50 μg/mL kanamycin were inoculated with 20 mL of an overnight culture of the transformed *E. coli* cells and incubated with shaking at 30 °C till reaching OD600 ~ 0.6. The overexpression of UvrD was induced by 0.5 mM Isopropyl β-D-1-thiogalactopyranoside (IPTG) concentration after which the culture was incubated with shaking at 27 °C for the period of 4 hrs. The cells were then isolated by centrifugation at 5,500x*g* for 10 min. The resulting pellet was resuspended into Lysis buffer (50 mM Tris.HCl pH 8, 500 mM NaCl, 40 mM Imidazole, 5 mM β-mercaptoethanol (BME), 5% glycerol and EDTA-free protease inhibitor cocktail tablet per 50 mL buffer). Subsequently, the cells were lysed by adding lysozyme to the final concentration of 2 mg/mL and kept at 4°C for 30 min followed by sonication. The crude lysate was then clarified by centrifugation at 95,000×*g*, for 1 hr at 4 °C. The supernatant was directly loaded onto HiTrap HP 5 ml affinity column (GE Healthcare) pre-equilibrated with Buffer A (50 mM Tris.HCl pH 8, 500 mM NaCl, 40 mM Imidazole, 5 mM BME and 5% glycerol). The column was washed with 50 ml of Buffer A afterwards the bound protein was eluted using linear gradient against Buffer B (50 mM Tris-HCl pH 8, 500 mM NaCl, 750 mM Imidazole, 5 mM BME and 5% glycerol). The eluted fractions containing His8-SUMO-UvrD were pooled and incubated with SUMO protease for 16 hrs at 4 °C to cleave the SUMO-tag and release native UvrD. After the digestion with SUMO protease, the solution was then loaded onto HiTrap HP 5 ml affinity column (GE Healthcare) pre-equilibrated with Buffer A. The flow-through fractions containing native UvrD were pooled and dialysed against storage buffer (50 mM Tris.HCl pH 8, 500 mM NaCl, 5 mM BME and 50% glycerol). The protein solution was further concentrated, flash frozen in liquid nitrogen, and stored at −80 °C. The protein concentration was calculated by measuring the absorbance at 280 nm and using the theoretical molecular extension coefficient estimated from the amino-acid sequence of UvrD (105770 M^-1^ cm^-1^).

### UvrD helicase reactions

UvrD helicase was a generous gift from Dr. Samir Hamdan. For neutravidin-displacement Substrate 1 or 2 (see Fig. 5a) was loaded into the flowcell (20 pM in Replicaiton Buffer supplemented with 150nM Sytox Orange) at a rate of 70 μL/min. For unwinding reactions Substrate 3 was loaded under the same conditions. After 1 minute or after a density of 0.3–0.7 molecule per μm^2^ on the surface was reached, the flowcell was washed with 140 μL of replication buffer at a rate of 70 μL/min. To remove excess DNA molecules from solution the flowcell was washed with 140 μL of replication buffer, for reactions on Substrate 1 or Substrate 2, supplemented with 100 nM UvrD. Finally, the reaction was initiated by loading the specified concentration of ATP in Replication buffer supplemented with 150nM Sytox Orange. Unwinding by UvrD was visualised with a constant buffer flow of 10 μL/min.

### Detection of events and post-synchronisation

Data analysis was carried out using Fiji [45] and Python. The raw data was first corrected for a non-uniform excitation-beam profile and mechanical drift of the microscope stage during the measurement (see supplementary Fig. 2,3). Next, all fluorescent spots corresponding to DNA templates bound to the surface were detected using a threshold approach (see supplementary Fig. 4) and the intensity of the DNA, and if present RPA, was measured over time. Next, all trajectories were fitted with the following piecewise linear function with three segments:

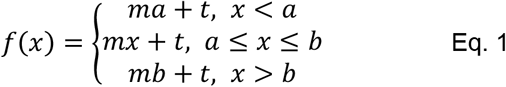

The parameter *a* denotes the time when enzymatic activity begins and the slope changes from 0 to a constant value *m.* The parameter *b* denotes the time when the whole substrate was processed and the slope becomes 0 again. The intensity during the first segment *(x<a)* corresponds to 2620 bp (or 0 bp in RPA trajectories), the last segment corresponds to 0 bp (or 2620 bp for RPA trajectories). The calibrated value *I_cal_* is then given by:

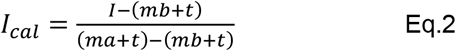

Where *I* denotes the raw fluorescence intensity. Again, for RPA trajectories increasing in intensity *a* has to be substituted with *b* and vice versa. Incomplete reactions, as determined by either negative parameters *a* or *b* or parameters *a* or *b* greater than the number of frames in a movie, were discarded. For Phi29 DNAp trajectories completion of replication was additionally confirmed by the dissociation of the newly synthesised double-stranded DNA from the now single-stranded template that remains bound to the cover slip surface (see Fig. 1a). This dissociation results in a sudden drop of intensity, detected by applying a regression tree algorithm [46], [47]. Trajectories considered for further analysis showed a coefficient of determination higher than 0.7. By defining the time of DNA dissociation for Phi29 trajectories or the parameter *b* for *λ exo* (see supplementary Fig 2) as time point zero, we synchronised trajectories at a well-defined time point corresponding to the end of the reaction (post-reaction synchronisation). Finally, we calculated the mean intensity in both channels at every time point, both before and after time point zero. Rate distributions for λ exo and UvrD were calculated by fitting the trajectories again to Eq.1, the absolute of fit parameter *m* then determines the rate.

### Determination of DNA synthesis rate by Phi29 DNAp

To determine the rate of DNA synthesis by Phi29 DNAp the post-synchronised trajectories were fit to to single exponential functions of the form:

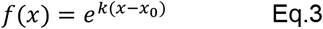

Where *k* denotes the rate constant in s^-1^, and *x_0_* shifts the function to negative time values. Values for *x>0*, i.e. after dissociation of the daughter strand where ignored for fits, since RPA fluorescence signal saturates and is no longer described by a single exponential function. To calculate a rate in bp·s^-1^ we multiplied the values with the length of the used substrate (2620 bp). To determine a Michaelis-Menten constant we plotted the obtained rates over the used dNTP concentration and fitted the data to the Michaelis-Menten equation:

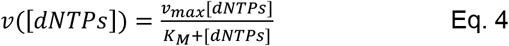

Where *v* denotes the synthesis rate as a function of dNTP concentration (*[dNTPs]*), *v_max_* denotes the synthesis rate in saturating dNTP concentrations and *K_M_* the Michaelis-Menten constant, the dNTP concentration at which the rate is half the saturation rate. Note that we assume an identical affinity for each of the four dNTPs.

### Determination of RPA association constant

We treated RPA binding to ssDNA as a first-order reaction. Such reactions are described by a differential equation of the form:

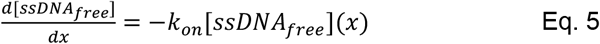

Where [ssDNA_free_] is the amount of free ssDNA that allows RPA binding and *k_on_* denotes the molecular rate association constant of RPA to ssDNA. By integration one finds:

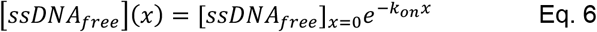

Where [ssDNA_free_]_x=0_ is the amount of initially available ssDNA. The total amount of ssDNA ([*ssDNA_tota_*]), free or bound cannot exceed the length of the template. It follows:

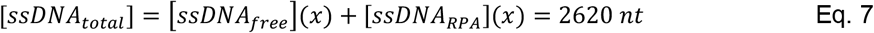

For the amount of RPA-bound ssDNA at any given time then follows:

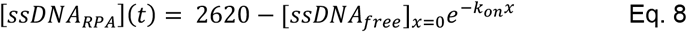

For fitting we introduced a time-offset x0 to account for the negative time values, by substituting *x* with (*x-x_0_*). This first order kintic only describes the reaction as the signal reaches saturation. We excluded any values for *x* < −20 s for the purpose of fitting.

### Determination of reaction intermediates in displacement of neutravidin by UvrD

To determine the number of intermediate steps in the reactions, we fit the data shown in Fig 5d to gamma-distributions of the form:

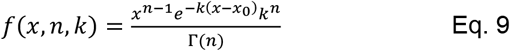

Eq. 9 The parameter *n* denotes the number of reaction intermediates with rate constant *k*. *Γ(n)* = *(n – 1)!* is the Gamma function, the value *x0* shifts the distribution along the x-axis and was fixed to 20 s for fitting, to account for the time when ATP reaches the flowcell and the reaction starts. Fitting was performed using the maxium likelihood estimation from the python package SciPy [48].

## DATA AVAILABILITY

Home-built ImageJ plugins and python scripts have been deposited on the Github repository for Single-molecule/Image analysis tools (https://github.com/Single-molecule-Biophysics-UOW).

## AUTHOR CONTRIBUTIONS

Conceptualisation, S.H.M., L.M.S., A.M.v.O.; Methodology, S.H.M., L.M.S., A.M.v.O.; Software, S.H.M.; Validation, S.H.M., L.J.F.; Formal analysis, S.H.M.; Investigation, S.H.M., L.J.F.; Resources, S.H.M., L.J.F., A.S.; Funding acquisition, L.M.S, S.M.H., A.M.v.O.; Supervision, L.M.S., S.M.H., A.M.v.O.; Visualisation, S.H.M.; Writing – original draft, S.H.M., L.M.S.,L.J.F., A.S.,S.M.H., A.M.v.O.

## ACKNOWLEDGEMENT

The authors thank Dr Jacob Lewis (University of Wollongong) and Prof. Michael O’Donnell (Rockefeller University) for contributing reagents.

This work was supported by the Australian Research Council (research grants DP150100956 and DP180100858 to A.M.v.O. and an Australian Laureate Fellowship FL140100027 to A.M.v.O.), the National Health and Medical Research Council (NHMRC Investigator grant 2007778 to L.M.S) and an Australian Government Research Training Program Scholarship (to S.H.M). Funding for open access charge: Australian Research Council.

## REFERENCES

[1] A. Robinson, R. J. Causer, and N. E. Dixon, “Architecture and Conservation of the Bacterial DNA Replication Machinery, an Underexploited Drug Target,” Curr. Drug Targets, vol. 13, no. 3, pp. 352–372, Feb. 2012, doi: 10.2174/138945012799424598.

[2] M. Macheret and T. D. Halazonetis, “DNA Replication Stress as a Hallmark of Cancer,” Annu. Rev. Pathol. Mech. Dis., vol. 10, no. 1, pp. 425–448, Jan. 2015, doi: 10.1146/annurev-pathol-012414-040424.

[3] P. S. Bernard and C. T. Wittwer, “Real-Time PCR Technology for Cancer Diagnostics,” Clin. Chem., vol. 48, no. 8, pp. 1178–1185, Aug. 2002, doi: 10.1093/clinchem/48.8.1178.

[4] I. Marota and F. Rollo, “Molecular paleontology,” Cellular and Molecular Life Sciences, vol. 59, no. 1. Cell Mol Life Sci, pp. 97–111, 2002, doi: 10.1007/s00018-002-8408-8.

[5] E. Monachino, L. M. Spenkelink, and A. M. van Oijen, “Watching cellular machinery in action, one molecule at a time.,” J. Cell Biol., vol. 216, no. 1, pp. 41–51, Jan. 2017, doi: 10.1083/jcb.201610025.

[6] J. S. Lewis et al., “Single-molecule visualization of fast polymerase turnover in the bacterial replisome,” Elife, 2017, doi: 10.7554/eLife.23932.

[7] J. S. Lewis et al., “Tunability of DNA Polymerase Stability during Eukaryotic DNA Replication,” Mol. Cell, Nov. 2019, Accessed: Dec. 03, 2019. [Online]. Available: https://www.sciencedirect.com/science/article/pii/S1097276519307658.

[8] L. M. Spenkelink et al., “Recycling of single-stranded DNA-binding protein by the bacterial replisome,” Nucleic Acids Res., vol. 47, no. 8, pp. 4111–4123, May 2019, doi: 10.1093/nar/gkz090.

[9] S. H. Mueller, L. M. Spenkelink, and A. M. van Oijen, “When proteins play tag: the dynamic nature of the replisome,” Biophys. Rev., 2019, doi: 10.1007/s12551-019-00569-4.

[10] M. R. Wasserman, G. D. Schauer, M. E. O’Donnell, and S. Liu, “Replication Fork Activation Is Enabled by a Single-Stranded DNA Gate in CMG Helicase,” Cell, vol. 178, no. 3, pp. 600–611.e16, Jul. 2019, doi: 10.1016/j.cell.2019.06.032.

[11] T. Suren, D. Rutz, P. Mößmer, U. Merkel, J. Buchner, and M. Rief, “Single-molecule force spectroscopy reveals folding steps associated with hormone binding and activation of the glucocorticoid receptor,” Proc. Natl. Acad. Sci. U. S. A., vol. 115, no. 46, pp. 11688–11693, Nov. 2018, doi: 10.1073/pnas.1807618115.

[12] A. M. van Oijen, P. C. Blainey, D. J. Crampton, C. C. Richardson, T. Ellenberger, and X. S. Xie, “Single-molecule kinetics of lambda exonuclease reveal base dependence and dynamic disorder.,” Science, vol. 301, no. 5637, pp. 1235–1238, Aug. 2003, doi: 10.1126/science.1084387.

[13] B. Ibarra et al., “Proofreading dynamics of a processive DNA polymerase,” EMBO J., vol. 28, no. 18, pp. 2794–2802, Sep. 2009, doi: 10.1038/emboj.2009.219.

[14] M. N. Dessinges, T. Lionnet, X. G. Xi, D. Bensimon, and V. Croquette, “Single-molecule assay reveals strand switching and enhanced processivity of UvrD,” Proc. Natl. Acad. Sci. U. S. A., vol. 101, no. 17, pp. 6439–6444, Apr. 2004, doi: 10.1073/PNAS.0306713101.

[15] K. Lee, H. Balci, H. Jia, T. M. Lohman, and T. Ha, “Direct imaging of single UvrD helicase dynamics on long single-stranded DNA,” Nat. Commun. 2013 41, vol. 4, no. 1, pp. 1–9, May 2013, doi: 10.1038/ncomms2882.

[16] R. Kovall and B. W. Matthews, “Toroidal structure of λ-exonuclease,” Science (80-.)., vol. 277, no. 5333, pp. 1824–1827, Sep. 1997, doi: 10.1126/SCIENCE.277.5333.1824/ASSET/8C1F719A-A79F-42B2-BC3E-0A4BDFAB840B/ASSETS/GRAPHIC/SE3875722004.JPEG.

[17] J. Zhang, K. A. McCabe, and C. E. Bella, “Crystal structures of λ exonuclease in complex with DNA suggest an electrostatic ratchet mechanism for processivity,” Proc. Natl. Acad. Sci. U. S. A., vol. 108, no. 29, pp. 11872–11877, Jul. 2011, doi: 10.1073/PNAS.1103467108/-/DCSUPPLEMENTAL.

[18] K. Subramanian, W. Rutvisuttinunt, W. Scott, and R. S. Myers, “The enzymatic basis of processivity in λ exonuclease,” Nucleic Acids Res., vol. 31, no. 6, pp. 1585–1596, Mar. 2003, doi: 10.1093/NAR/GKG266.

[19] S. I. Matsuura et al., “Real-time observation of a single DNA digestion by λ exonuclease under a fluorescence microscope field,” Nucleic Acids Res., vol. 29, no. 16, pp. e79–e79, Aug. 2001, doi: 10.1093/NAR/29.16.E79.

[20] P. H. Lu, L. Xun, and S. X. Xie, “Single-Molecule Enzymatic Dynamics,” Science (80-.)., vol. 282, no. 5395, pp. 1877–1882, Dec. 1998, doi: 10.1126/SCIENCE.282.5395.1877.

[21] J. S. Lewis et al., “Single-molecule visualization of Saccharomyces cerevisiae leading-strand synthesis reveals dynamic interaction between MTC and the replisome,” Proc. Natl. Acad. Sci., vol. 114, no. 40, pp. 10630–10635, 2017, doi: 10.1073/pnas.1711291114.

[22] B. Gibb et al., “Concentration-Dependent Exchange of Replication Protein A on Single-Stranded DNA Revealed by Single-Molecule Imaging,” PLoS One, vol. 9, no. 2, p. e87922, Feb. 2014, doi: 10.1371/journal.pone.0087922.

[23] N. Pokhrel et al., “Monitoring Replication Protein A (RPA) dynamics in homologous recombination through site-specific incorporation of non-canonical amino acids,” Nucleic Acids Res., vol. 45, no. 16, pp. 9413–9426, Sep. 2017, doi: 10.1093/NAR/GKX598.

[24] J. Chen, S. Le, A. Basu, W. J. Chazin, and J. Yan, “Mechanochemical regulations of RPA’s binding to ssDNA,” Sci. Rep., vol. 5, no. 1, p. 9296, Aug. 2015, doi: 10.1038/srep09296.

[25] C. Veigel, F. Wang, M. L. Bartoo, J. R. Sellers, and J. E. Molloy, “The gated gait of the processive molecular motor, myosin V,” Nat. Cell Biol., vol. 4, no. 1, pp. 59–65, Dec. 2002, doi: 10.1038/ncb732.

[26] C. Chen et al., “Single-Molecule Fluorescence Measurements of Ribosomal Translocation Dynamics,” Mol. Cell, vol. 42, no. 3, pp. 367–377, May 2011, doi: 10.1016/j.molcel.2011.03.024.

[27] J. A. Morin et al., “Mechano-chemical kinetics of DNA replication: identification of the translocation step of a replicative DNA polymerase,” Nucleic Acids Res., vol. 43, no. 7, p. 3643, Feb. 2015, doi: 10.1093/NAR/GKV204.

[28] F. E. Kemmerich, P. Daldrop, C. Pinto, M. Levikova, P. Cejka, and R. Seidel, “Force regulated dynamics of RPA on a DNA fork,” Nucleic Acids Res., vol. 44, no. 12, pp. 5837–5848, Jul. 2016, doi: 10.1093/nar/gkw187.

[29] V. Petrova et al., “Active displacement of RecA filaments by UvrD translocase activity,” Nucleic Acids Res., vol. 43, no. 8, p. 4133, Feb. 2015, doi: 10.1093/NAR/GKV186.

[30] N. K. Maluf, C. J. Fischer, and T. M. Lohman, “A Dimer of Escherichia coli UvrD is the Active Form of the Helicase In Vitro,” J. Mol. Biol., vol. 325, no. 5, pp. 913–935, Jan. 2003, doi: 10.1016/S0022-2836(02)01277-9.

[31] M. C. Hall, J. R. Jordan, and S. W. matson, “Evidence for a physical interaction between the Escherichia coli methyl-directed mismatch repair proteins MutL and UvrD,” EMBO J., vol. 17, no. 5, pp. 1535–1541, Mar. 1998, doi: 10.1093/EMBOJ/17.5.1535.

[32] S. P. Carney et al., “Kinetic and structural mechanism for DNA unwinding by a non-hexameric helicase,” Nat. Commun. 2021 121, vol. 12, no. 1, pp. 1–14, Dec. 2021, doi: 10.1038/s41467-021-27304-6.

[33] E. Curti, S. J. Smerdon, and E. O. Davis, “Characterization of the Helicase Activity and Substrate Specificity of Mycobacterium tuberculosis UvrD,” J. Bacteriol., vol. 189, no. 5, p. 1542, Mar. 2007, doi: 10.1128/JB.01421-06.

[34] H. W. McKenzie, M. A. Lewis, and E. H. Merrill, “First passage time analysis of animal movement and insights into the functional response,” Bull. Math. Biol., vol. 71, no. 1, pp. 107–129, Jan. 2009, doi: 10.1007/S11538-008-9354-X.

[35] A. K. Verma, A. Bhatnagar, D. Mitra, and R. Pandit, “First-passage-time problem for tracers in turbulent flows applied to virus spreading,” Phys. Rev. Res., vol. 2, no. 3, p. 033239, Aug. 2020, doi: 10.1103/PHYSREVRESEARCH.2.033239/FIGURES/4/MEDIUM.

[36] D. R. Burnham, H. B. Kose, R. B. Hoyle, and H. Yardimci, “The mechanism of DNA unwinding by the eukaryotic replicative helicase,” Nat. Commun. 2019 101, vol. 10, no. 1, pp. 1–14, May 2019, doi: 10.1038/s41467-019-09896-2.

[37] D. L. Floyd, S. C. Harrison, and A. M. Van Oijen, “Analysis of Kinetic Intermediates in SingleParticle Dwell-Time Distributions,” Biophys. J., vol. 99, no. 2, pp. 360–366, Jul. 2010, doi: 10.1016/J.BPJ.2010.04.049.

[38] J. A. Ali and T. M. Lohman, “Kinetic measurement of the step size of DNA unwinding by Escherichia coli UvrD helicase,” Science (80-.)., vol. 275, no. 5298, pp. 377–380, Jan. 1997, doi: 10.1126/science.275.5298.377.

[39] B. Nguyen, Y. Ordabayev, J. E. Sokoloski, E. Weiland, and T. M. Lohman, “Large domain movements upon UvrD dimerization and helicase activation,” Proc. Natl. Acad. Sci. U. S. A., vol. 114, no. 46, pp. 12178–12183, Nov. 2017, doi: 10.1073/PNAS.1712882114/-/DCSUPPLEMENTAL.

[40] T. A. Kunkel and D. A. Erie, “DNA Mismatch Repair,” Annu. Rev. Biochem., vol. 74, no. 1, pp. 681–710, Jun. 2005, doi: 10.1146/annurev.biochem.74.082803.133243.

[41] P. Montero L. et al., “Best practices and tools for reporting reproducible fluorescence microscopy methods,” Nat. Methods, vol. 18, no. 12, pp. 1463–1476, Dec. 2021, doi: 10.1038/S41592-021-01156-W.

[42] S. H. Mueller, L. M. Spenkelink, A. M. Van Oijen, and J. S. Lewis, “Design of customizable long linear DNA substrates with controlled end modifications for single-molecule studies,” Anal. Biochem., vol. 592, p. 113541, Mar. 2020, doi: 10.1016/j.ab.2019.113541.

[43] A. O. HJ Geertsema, KE Duderstadt, “Single-Molecule Observation of Prokaryotic DNA Replication,” Methods Mol Biol, vol. 1300, pp. 219–238, 2015, doi: 10.1007/978-1-4939-2596-4_14.

[44] L. D. Langston et al., “CMG helicase and DNA polymerase form a functional 15-subunit holoenzyme for eukaryotic leading-strand DNA replication,” Proc. Natl. Acad. Sci., vol. 111, no. 43, pp. 15390–15395, Oct. 2014, doi: 10.1073/pnas.1418334111.

[45] J. Schindelin et al., “Fiji: an open-source platform for biological-image analysis,” Nat. Methods, vol. 9, no. 7, pp. 676–682, Jul. 2012, doi: 10.1038/nmeth.2019.

[46] L. Breiman, J. Friedman, C. J. Stone, and R. A. Olshen, Classification and Regression Trees. 1984.

[47] F. Pedregosa et al., “Scikit-learn: Machine Learning in Python,” 2011. Accessed: Feb. 08, 2021. [Online]. Available: http://scikit-learn.sourceforge.net.

[48] P. Virtanen et al., “SciPy 1.0: fundamental algorithms for scientific computing in Python,” Nat. Methods, vol. 17, no. 3, pp. 261–272, Mar. 2020, doi: 10.1038/S41592-019-0686-2.

