## Supplementary Information for "Rapid single-molecule characterisation of nucleic-acid enzymes"

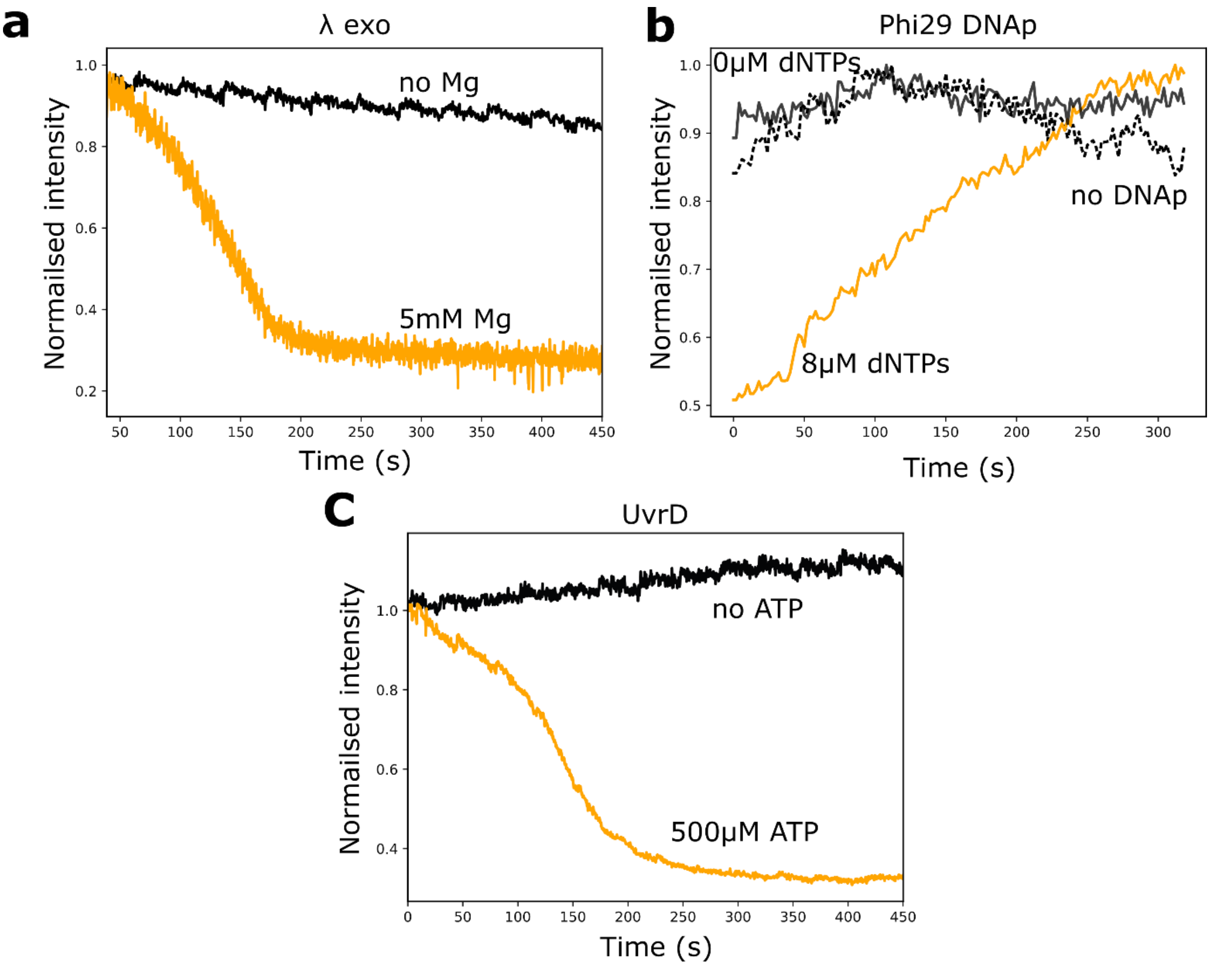


***Supplementary Fig. 1: Control reactions. (a)*** *To confirm the observed signal is due to λ exo activity we imaged in absence of Mg but otherwise identical conditions as stated in methods. This control also shows that photo-bleaching does not affect our quantification of dsDNA, since S.O. signal remains approximately constant over whole observed time scale in the absence of enzymatic activity. The shown trajectories are non-synchronised averages, orange line: no Mg, black line: with Mg.* ***(b)*** *non-synchronised averages of Phi29 DNAp trajectories (see methods) with 8 μM dNTPs (orange), 0 μM dNTPs (solid black line) and 8 μM dNTPs but in absence of Phi29 DNAp (dashed black line). The black lines also prove that our observed RPA kinetics are not influenced by photo-bleaching. They remain approximately constant in the absence of enzymatic activity.* ***(c)*** *UvrD on substrate 3 (see methods) in presence of 500 μM ATP (orange) and in absence of ATP (black), with a buffer flow of 10 μL min*$^{-1}$.

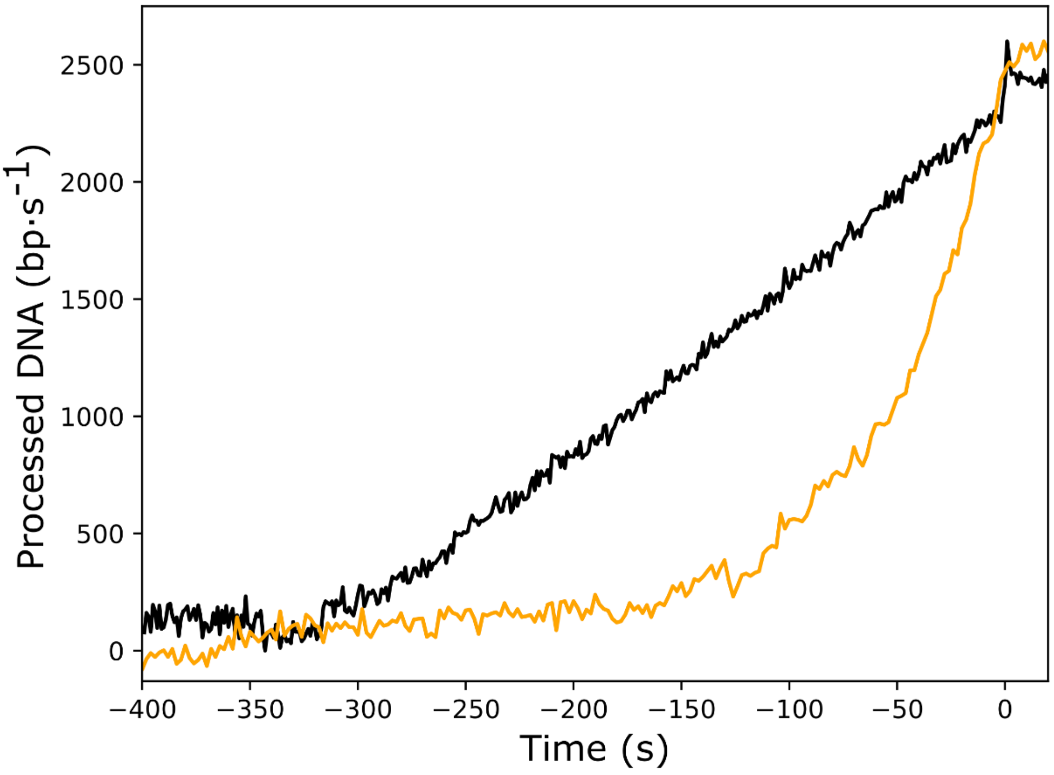


***Supplementary Fig. 2: Post-synchronised average trajectories.*** *To prove that the non-linear behaviour in Phi29 DNAp averages is not due to synchronisation, we synchronised λ exo trajectories, determining the end point of individual trajectories using a piecewise linear fit (see methods). The graph shows the inverted synchronised mean of 666 λ exo trajectories (black) to provide easy comparison to the synchronised mean of 127 Phi29 DNAp trajectories (orange).*

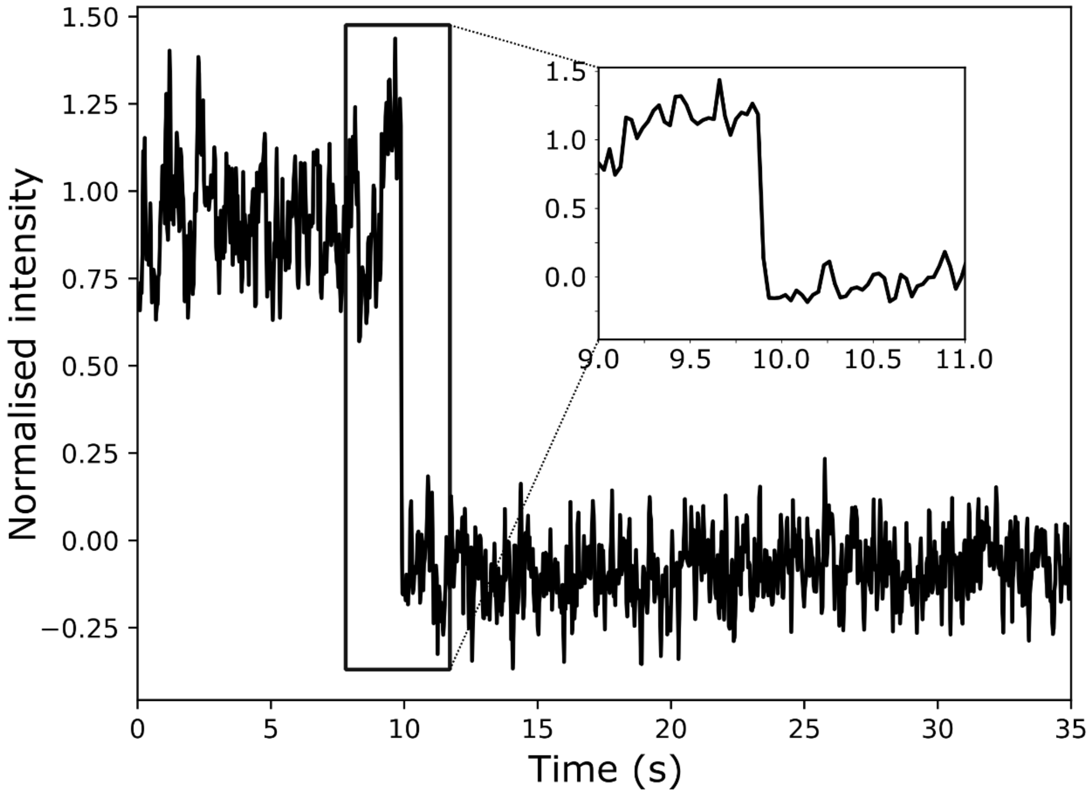


***Supplementary Fig. 3: High-Speed imaging of UvrD dependent dissociation of substrate 1.*** *Substrate 1, stained by S.O. was imaged in presence of 100 nM UvrD and 10 µM ATP with an exposure time of 30 ms and a laser intensity of 59 mW·cm². The laser intensity drops within one frame, i.e. 30 ms.*

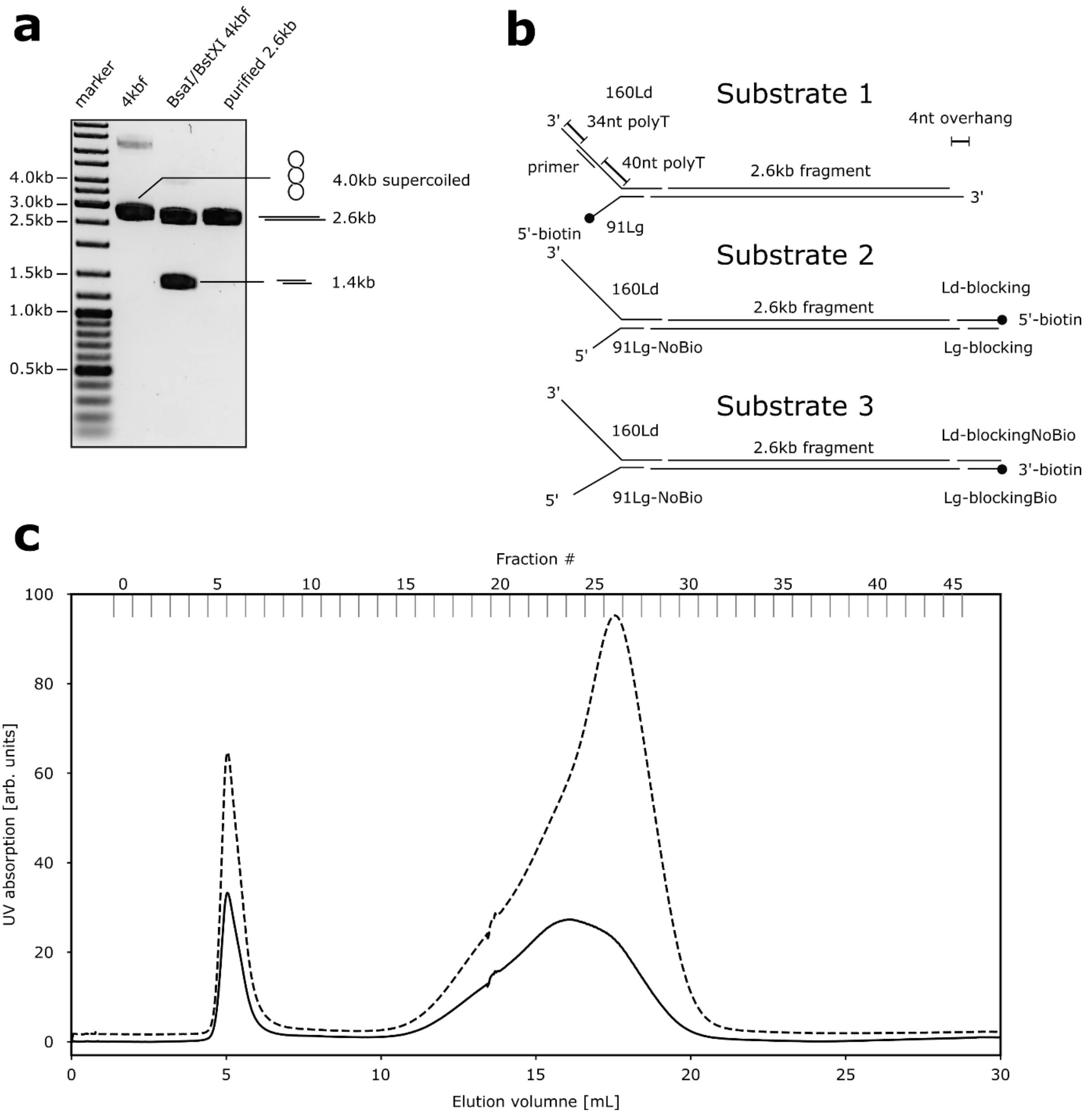


***Supplementary Fig. 4: Construction of 2.6kb DNA. (A)*** *1% agarose gel, staining with 0.5 ug/mL EtBr. Lane 2 shows the supercoiled 4kbf plasmid, Lane 3 the digested plasmid including the 2.6kb fragment. The 2.6kb fragment was excised from a separate gel and is shown in lane 4. After ligation of blocking and fork oligonucleotides (see **B** and supplementary table 1) the final template was purified on a Sepharose 4B column. **(C)** Size-exclusion chromatogram showing UV280 absorption (solid line) and UV260 absorption (dashed line). The first peak corresponds to the 2.6kb DNA, the second peak consists of excess oligonucleotides and T4 ligase.*

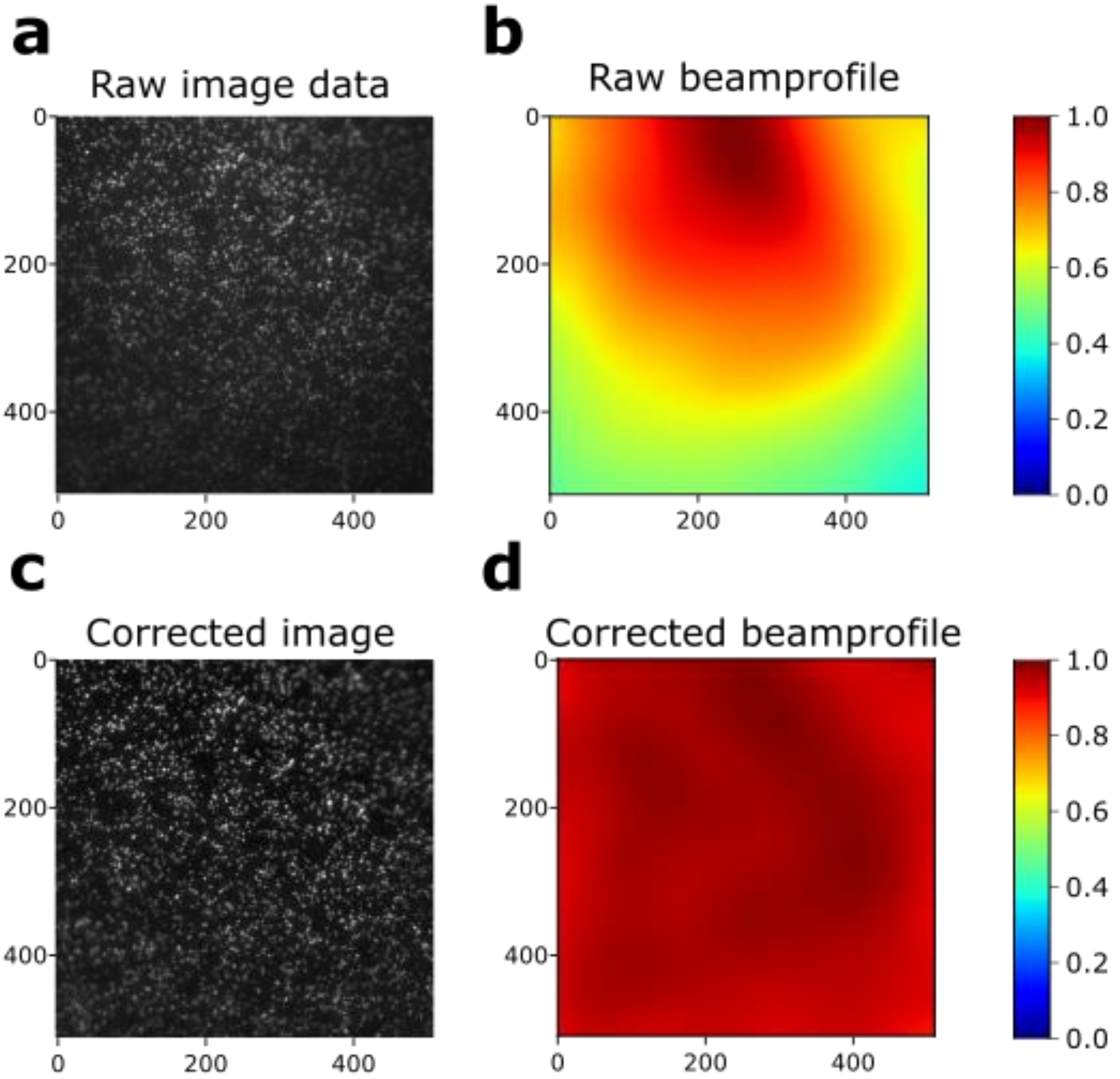


***Supplementary Fig. 5: Beam profile correction:*** *Our assay relies on reproducible and quantifiable fluorescence intensity measurement. However, uneven illumination of samples can lead to differences in fluorescence intensity within single images.* ***(A)*** *shows raw microscopic image data. Note that spots near the edges are considerably dimmer than spots close to the middle.* ***(B)*** *Applying a gaussian blur to the image evens out local intensity differences and yields the beam profile, here shown as a heatmap.* ***(C)*** *The raw image is corrected by dividing every pixel value by the beam profile.* ***(D)*** *The quality of the correction can be assessed by applying a gaussian blur once again. Comparing this beam profile to the raw beam profile (see (B)) one can see that large scale intensity fluctuations are now largely eliminated. To assure reusability of our method we implemented this correction in the form of an ImageJ plugin. We utilize the imglib2 java library*[1] *to achieve generic processing of single images, movies or multi-colour movies alike. See https://github.com/Single-molecule-Biophysics-UOW for source code and download.*

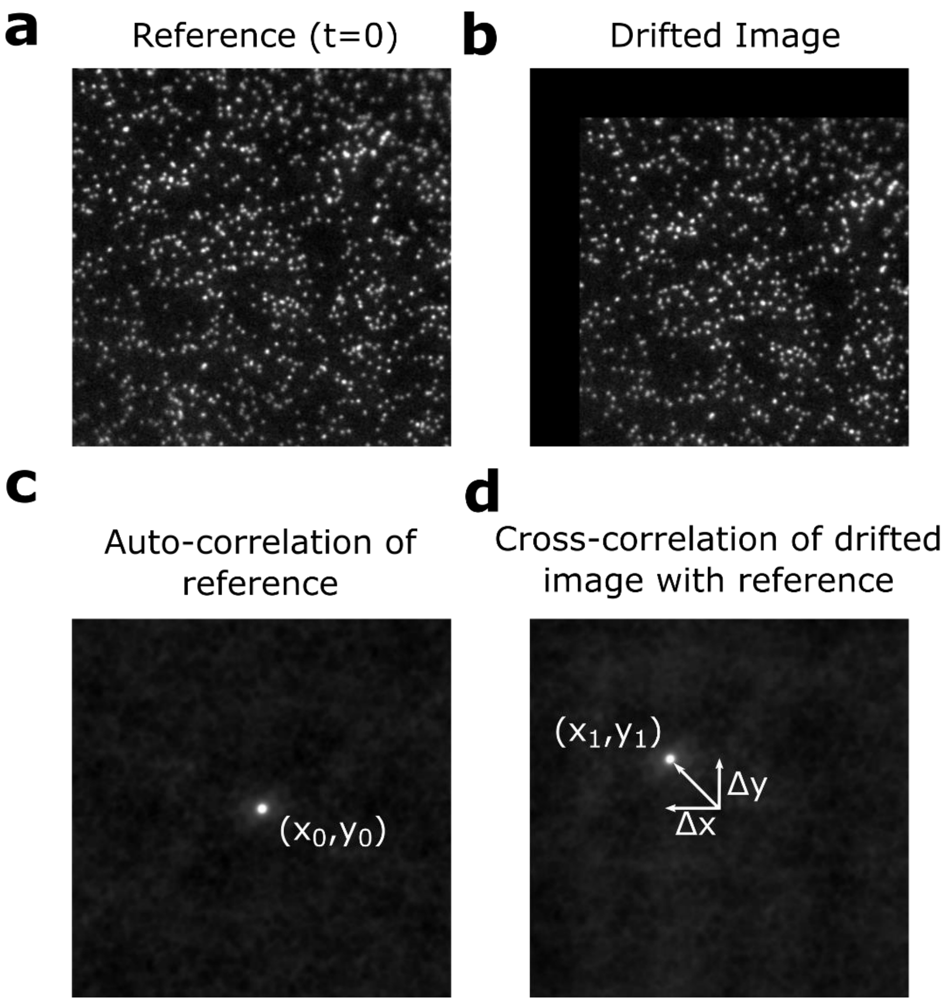


***Supplementary Fig. 6: Drift correction.*** *To correct for mechanical drift of the imaged sample we implement a template based correction algorithm. Briefly, the first frame of a movie* ***(a)*** *is treated as the reference.* ***(b)*** *shows the reference shifted in x and y direction to simulate drift. To find the total shift the cross-correlation* ***(d)*** *of the image with the reference is compared to the auto-correlation of the reference* ***(c)****. The shift-vector (Δx,Δy) is given by the change of position of the maximum value of the correlation images. For fast image-processing correlations are calculated in the Fourier-domain. We utilize the imglib2 java library* [1] *to achieve generic processing of single images, movies or multi-colour movies alike. See [https://github.com/Single-molecule-Biophysics-UOW](https://github.com/Single-molecule-Biophysics-UOW) for source code and download.*

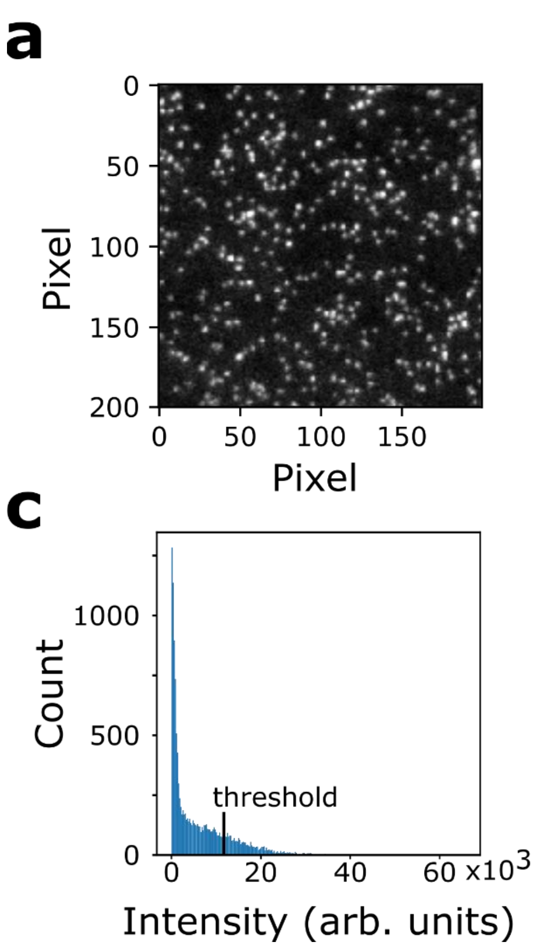


***Supplementary Fig. 7: Detection of events.*** *As described in methods section, we use a thresholding approach to detect the signal of single DNA molecules.* ***(a)*** *shows a 200x200 pixel region from supplementary Fig 2c. Before threshold analysis a discoidal filter is applied.* ***(b)*** *Filtered image. The discoidal filter is a moving window filter. Every pixel's grey value is substituted by the mean of all pixels within an inner radius $r_{in}$.* Subsequently the mean grey value of all pixels within a second outer region with radius $r_{out}$ is subtracted. For the shown image $r_{in}$ =1 and $r_{out}$=3. This leads to greatly reduces background intensity. **(c)** *histogram of grey values in* ***(b).*** *To detect peaks a threshold is set to be twice the standard deviation of the distribution of grey values, as indicated by the black bar.* ***(d)*** *detected peaks. Every pixel with a grey value above the threshold is defined as one peak. If multiple pixel coincide within a minimum distance of 6 pixels only the one with the higher grey value is counted as peak.*

*Table 7.1: Sequences of oligonucleotides used in this study.*

| **Name (DNA)** | **Sequence (5'—3')** |
|---|---|
| BlockingLdNoBio | AGT CGC AGC TAT AGG TGG CAT TTC AG |
| BlockingLd | /Bio/AGT CGC AGC TAT AGG TGG CAT TTC AG |
| BlockingLg | /Phos/CTG AAA TGC CAC CTA TAG CTG CGA CTC ATG |
| BlockingLgBio | /Phos/CTG AAA TGC CAC CTA TAG CTG CGA CTC ATG/Bio/ |
| 160Ld | /Phos/ACC GAT GTG GTA GGA AGT GAG AAT TGG AGA GTG TGT TTT TTT TTT TTT TTT TTT TTT TTT TTT TTT TTT TTT TTT GAG GAA AGA ATG TTG GTG AGG GTT GGG AAG TGG AAG GAT GGG CTC GAG AGG TTT TTT TTT TTT TTT TTT TTT TTT TTT TTT T*T*T*T |
| 91Lg | TTT TTT TTT TTT TTT TTT TTT TTT TTT TTT TTT TTT TTT TTT TTT TTT TTT TTT TTT TTT CAC ACT CTC CAA TTC TCA CTT CCT ACC ACA T |
| Fork Primer | CCT CTC GAG CCC ATC CTT CCA CTT CCC AAC CCT CAC C |

*Supplementary Table 2: Complete oligonucleotide sequence of 4kbf.*

ACCGCGAG ACCCACGC TCACCGGC TCCAGATT TATCAGCA ATAAACCA GCCAGCCG
GAAGGGCC GAGCGCAG AAGTGGTC CTGCAACT TTATCCGC CTCCATCC AGTCTATT
AATTGTTG CCGGGAAG CTAGAGTA AGTAGTTC GCCAGTTA ATAGTTTG CGCAACGT
TGTTGCCA TTGCTACA GGCATCGT GGTGTCAC GCTCGTCG TTTGGTAT GGCTTCAT
TCAGCTCC GGTTCCCA ACGATCAA GGCGAGTT ACATGATC CCCCATGT TGTGCAAA
AAAGCGGT TAGCTCCT TCGGTCCT CCGATCGT TGTCAGAA GTAAGTTG GCCGCAGT
GTTATCAC TCATGGTT ATGGCAGC ACTGCATA ATTCTCTT ACTGTCAT GCCATCCG
TAAGATGC TTTTCTGT GACTGGTG AGTACTCA ACCAAGTC ATTCTGAG AATAGTGT
ATGCGGCG ACCGAGTT GCTCTTGC CCGGCGTC AATACGGG ATAATACC GCGCCACA
TAGCAGAA CTTTAAAA GTGCTCAT CATTGGAA AACGTTCT TCGGGGCG AAAACTCT
CAAGGATC TTACCGCT GTTGAGAT CCAGTTCG ATGTAACC CACTCGTG CACCCAAC
TGATCTTC AGCATCTT TTACTTTC ACCAGCGT TTCTGGGT GAGCAAAA ACAGGAAG
GCAAAATG CCGCAAAA AAGGGAAT AAGGGCGA CACGGAAA TGTTGAAT ACTCATAC
TCTTCCTT TTTCAATA TTATTGAA GCATTTAT CAGGGTTA TTGTCTCA TGAGCGGA
TACATATT TGAATGTA TTTAGAAA AATAAACA AATAGGGG TTCCGCGC ACATTTCC
CCGAAAAG TGCCACCT GACGTCTA AGAAACCA TTATTATC ATGACATT AACCTATA
AAAATAGG CGTATCAC GAGGCCCT TTCGTCTC GCGCGTTT CGGTGATG ACGGTGAA
AACCTCTG ACACATGC AGCTCCCG GAGACGGT CACAGCTT GTCTGTAA GCGGATGC
CGGGAGCA GACAAGCC CGTCAGGG CGCGTCAG CGGGTGTT GGCGGGTG TCGGGGCT
GGCTTAAC TATGCGGC ATCAGAGC AGATTGTA CTGAGAGT GCACCATA TGCGGTGT
GAAATACC GCACAGAT GCGTAAGG AGAAAATA CCGCATCA GGCGCCAT TCGCCATT
CAGGCTGC GCAACTGT TGGGAAGG GCGATCGG TGCGGGCC TCTTCGCT ATTACGCC
AGCTGGCG AAAGGGGG ATGTGCTG CAAGGCGA TTAAGTTG GGTAACGC CAGGGTTT
TCCCAGTC ACGACGTT GTAAAACG ACGGCCAG TGAATTCG AGCTCGGA TGTTTTGG
CTCTGGTC AATGATTA CGGCATTG ATATCGTC CAACTGCA TGGAGATG AGTCGTGG
CAAGAATA CCAAGAGT TCCTCGGT TTGCCAGT TATTAAAA GACTCGTA TTTCCAAA
AGACTGCA ACATACTA CTCAGTGC AGCTTCAC AGAAACCT CATTCGTT TATTCCCT
TGTTTGAT TCAGAAGC AGGTGGGA CAGGTGAA CTTTTGGA TTGGAACT CGATTTCT
GACTGGGT TGGAAGGC AAGAGAGC CCCGAAAG CTTACATT TTATGTTA GCTGGTGG
ACTGACGC CAGAAAAT GTTGGTGA TGCGCTTA GATTAAAT GGCGTTAT TGGTGTTG
ATGTAAGC GGAGGTGT GGAGACAA ATGGTGTA AAAGACTC TAACAAAA TAGCAAAT
TTCGTCAA AAATGCTA AGAAATAG GTTATTAC TGAGTAGT ATTTATTT AAGTATTG
TTTGTGCA CTTGCCTG CAGGCCTT TTGAAAAG CAAGCATA AAAGATCT AAACATAA
AATCTGTA AAATAACA AGATGTAA AGATAATG CTAAATCA TTTGGCTT TTTGATTG
ATTGTACA GGAAAATA TACATCGC AGGGGGTT GACTTTTA CCATTTCA CCGCAATG
GAATCAAA CTTGTTGA AGAGAATG TTCACAGG CGCATACG CTACAATG ACCCGATT
CTTGCTAG CCTTTTCT CGGTCTTG CAAACAAC CGCCGGCA GCTTAGTA TATAAATA
CACATGTA CATACCTC TCTCCGTA TCCTCGTA ATCATTTT CTTGTATT TATCGTCT
TTTCGCTG TAAAAACT TTATCACA CTTATCTC AAATACAC TTATTAAC CGCTTTTA
CTATTATC TTCTACGC TGACAGTA ATATCAAA CAGTGACA CATATTAA ACACAGTG
GTTTCTTT GCATAAAC ACCATCAG CCTCAAGT CGTCAAGT AAAGATTT CGTGTTCA
TGCAGATA GATAACAA TCTATATG TTGATAAT TAGCGTTG CCTCATCA ATGCGAGA
TCCGTTTA ACCGGACC CTAGTGCA CTTACCCC ACGTTCGG TCCACTGT GTGCCGAA
CATGCTCC TTCACTAT TTTAACAT GTGGAATT CTTGAAAG AATGAGTT CAGTGGTG
CTGACACG ATATTTTA TTCATACA ACATTGAT TTCACCAA TTATAATA CCGCGCAA
TACTGTAT GAGCATAC AGTGGATC CGAGTCAT TCCTGCAG CGAGTCCA TGGGAGTC
AAATAGAC AACGATTT GAATTCGC TCTTCCGC TCGCGGCC GCTAAGAC TCGAGTAG
ATGACTAC GAGGTACC CGGGGATC CTCTAGAG TCGACCTG CAGGCATG CAAGCTTG
GCGTAATC ATGGTCAT AGCTGTTT CCTGTGTG AAATTGTT ATCCGCTC ACAATTCC
ACACAACA TACGAGCC GGAAGCAT AAAGTGTA AAGCCTGG GGTGCCTA ATGAGTGA

GCTAACTC ACATTAAT TGCGTTGC GCTCACTG CCCGCTTT CCAGTCGG GAAACCTG
TCGTGCCA GCTGCATT AATGAATC GGCCAACG CGCGGGGA GAGGCGGT TTGCGTAT
TGGGCGCT CTTCCGCT TCCTCGCT CACTGACT CGCTGCGC TCGGTCGT TCGGCTGC
GGCGAGCG GTATCAGC TCACTCAA AGGCGGTA ATACGGTT ATCCACAG AATCAGGG
GATAACGC AGGAAAGA ACATGTGA GCAAAAGG CCAGCAAA AGGCCAGG AACCGTAA
AAAGGCCG CGTTGCTG GCGTTTTT CCATAGGC TCCGCCCC CCTGACGA GCATCACA
AAAATCGA CGCTCAAG TCAGAGGT GGCGAAAC CCGACAGG ACTATAAA GATACCAG
GCGTTTCC CCCTGGAA GCTCCCTC GTGCGCTC TCCTGTTC CGACCCTG CCGCTTAC
CGGATACC TGTCCGCC TTTCTCCC TTCGGGAA GCGTGGCG CTTTCTCA TAGCTCAC
GCTGTAGG TATCTCAG TTCGGTGT AGGTCGTT CGCTCCAA GCTGGGCT GTGTGCAC
GAACCCCC CGTTCAGC CCGACCGC TGCGCCTT ATCCGGTA ACTATCGT CTTGAGTC
CAACCCGG TAAGACAC GACTTATC GCCACTGG CAGCAGCC ACTGGTAA CAGGATTA
GCAGAGCG AGGTATGT AGGCGGTG CTACAGAG TTCTTGAA GTGGTGGC CTAACTAC
GGCTACAC TAGAAGAA CAGTATTT GGTATCTG CGCTCTGC TGAAGCCA GTTACCTT
CGGAAAAA GAGTTGGT AGCTCTTG ATCCGGCA AACAAACC ACCGCTGG TAGCGGTG
GTTTTTTT GTTTGCAA GCAGCAGA TTACGCGC AGAAAAAA AGGATCTC AAGAAGAT
CCTTTGAT CTTTTCTA CGGGGTCT GACGCTCA GTGGAACG AAAACTCA CGTTAAGG
GATTTTGG TCATGAGA TTATCAAA AAGGATCT TCACCTAG ATCCTTTT AAATTAAA
AATGAAGT TTTAAATC AATCTAAA GTATATAT GAGTAAAC TTGGTCTG ACAGTTAC
CAATGCTT AATCAGTG AGGCACCT ATCTCAGC GATCTGTC TATTTCGT TCATCCAT
AGTTGCCT GACTCCCC GTCGTGTA GATAACTA CGATACGG GAGGGCTT ACCATCTG
GCCCCAGT GCTGCAAT GAT
